## Supplementary Information for "Hippo signaling differentially regulates distal progenitor subpopulations and their transitional states to construct the mammalian lungs"

Zhang et al. Supplementary Figure 1

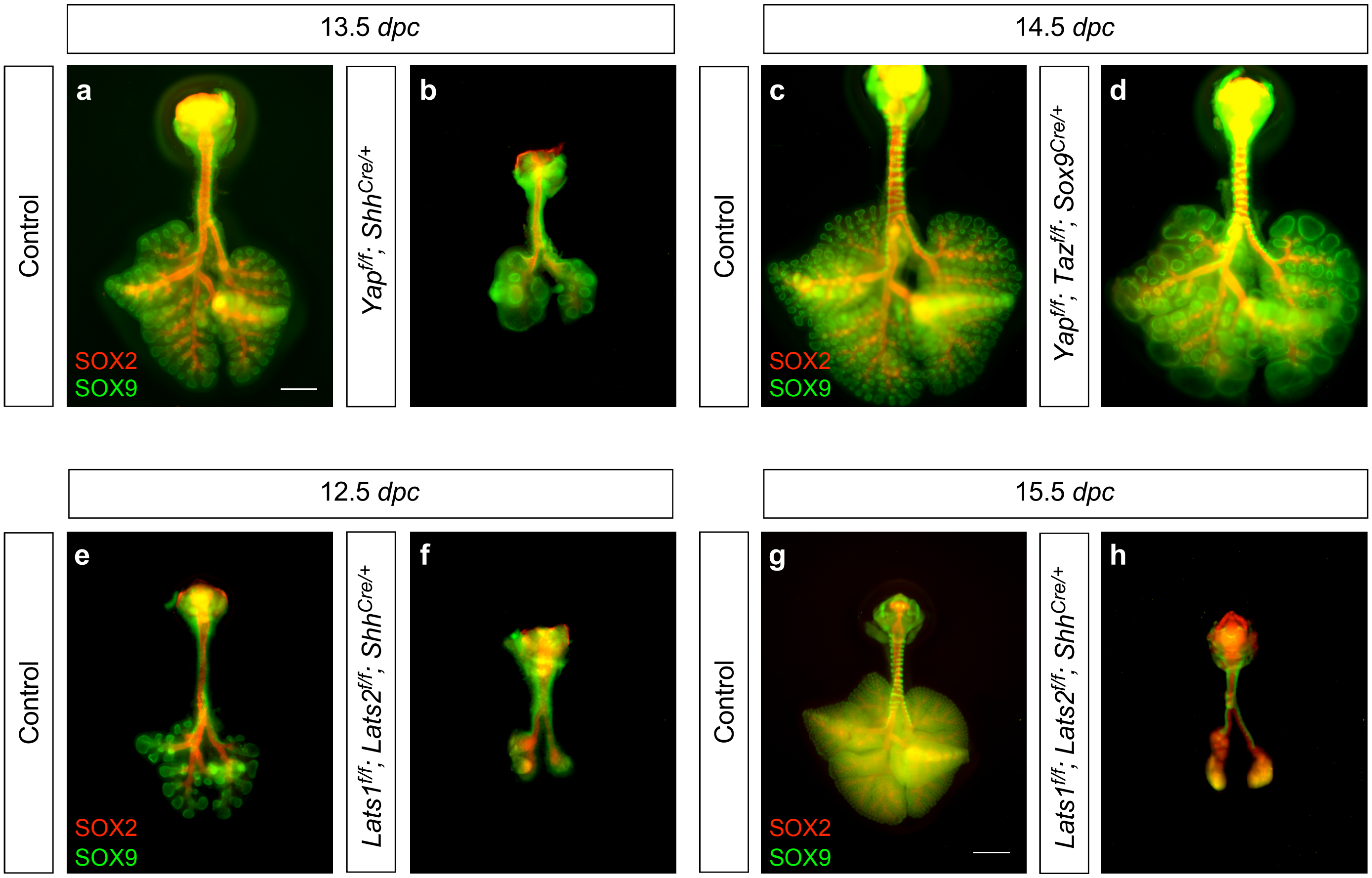

**Supplementary Fig. 1 The SOX9$^+$ domain is reduced when the Hippo pathway is perturbed**

(a, b) Whole-mount immunostaining of dissected lungs (ventral view) from control and *Yap$^{f/f}$; Shh$^{Cre/+}$* mice at 13.5 *days post-coitus* (*dpc*). *Shh$^{Cre}$* converted a floxed allele of *Yap* (*Yap$^f$*) into a null allele. SOX9 and SOX2 marked the distal and proximal airway epithelium, respectively. A reduction in SOX9 and SOX2 signals was discerned when *Yap* was eliminated. The *Shh$^{Cre}$* allele disrupted the *Shh* gene and could contribute to the observed phenotype. (c, d) Whole-mount immunostaining of dissected lungs from control and *Yap$^{f/f}$; Taz$^{f/f}$; Sox9$^{Cre/+}$* (Mutant) mice at 14.5 *dpc*. *Sox9$^{Cre}$* converted floxed alleles of *Yap* (*Yap$^f$*) and *Taz* (*Taz$^f$*) into null alleles. *Taz* is a minor player in lung branching, and its removal did not exacerbate the lung defects resulting from *Yap* loss. Distal lung cysts were present in the mutant lungs, and the size of the SOX9$^+$ pool appeared to be reduced. Quantitative analysis is required to accurately assess the changes in the SOX9$^+$ pool size. (e, f) Whole-mount immunostaining of dissected lungs from control and *Lats1$^{f/f}$; Lats2$^{f/f}$; Shh$^{Cre/+}$* mice at 12.5 *dpc*. Both the SOX2$^+$ and SOX9$^+$ domains were severely diminished. (g, h) Whole-mount immunostaining of dissected lungs from control and *Lats1$^{f/f}$; Lats2$^{f/f}$; Shh$^{Cre/+}$* mice at 15.5 *dpc*. The size reduction of both the SOX2$^+$ and SOX9$^+$ domains persisted as lung development proceeded.

Scale bars, 0.5 mm (a-f), 1 mm (g, h).

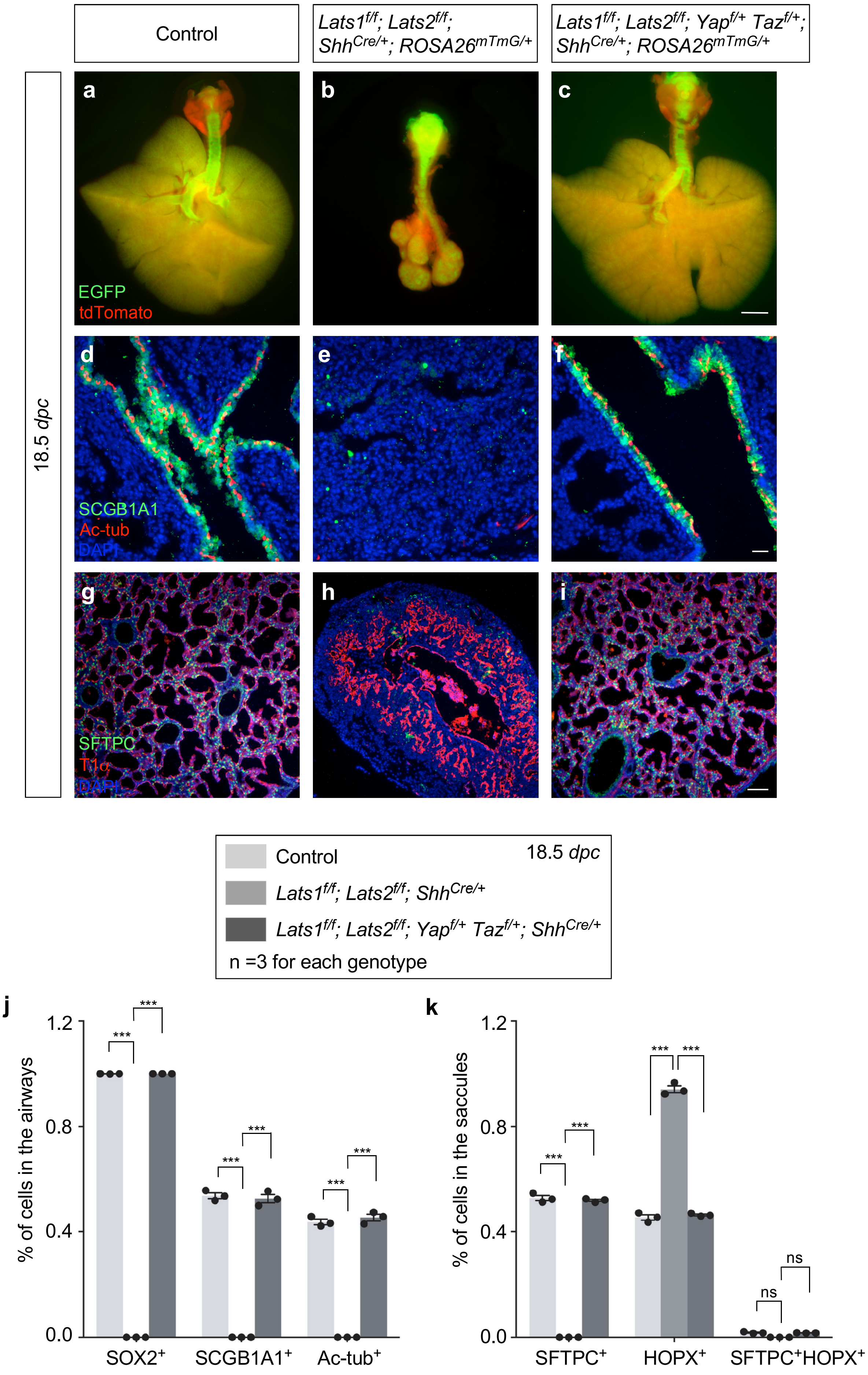

Control
Lats1f/f; Lats2f/f; ShhCre/+; ROSA26mTmG/+
Lats1f/f; Lats2f/f; Yapf/+ Tazf/+; ShhCre/+; ROSA26mTmG/+
18.5 dpc
a
b
c
EGFP
tdTomato
d
e
f
SCGB1A1
Ac-tub
DAPI
g
h
i
SFTPC
T1α
DAPI
Control
18.5 dpc
Lats1f/f; Lats2f/f; ShhCre/+
Lats1f/f; Lats2f/f; Yapf/+ Tazf/+; ShhCre/+
n =3 for each genotype
j
% of cells in the airways
1.2
0.8
0.4
0.0
***
SOX2+
SCGB1A1+
Ac-tub+
k
% of cells in the saccules
ns
SFTPC+
HOPX+
SFTPC+HOPX+

**Supplementary Fig. 2 Loss of one copy of *Yap* and *Taz* partially rescues the lung defects in *Lats1$^{f/f}$; Lats2$^{f/f}$; Shh$^{Cre/+}$* mice**

(a-c) Whole-mount imaging of lungs (ventral view) from control, *Lats1$^{f/f}$; Lats2$^{f/f}$; Shh$^{Cre/+}$; ROSA26$^{mTmG/+}$* (Mutant) and *Lats1$^{f/f}$; Lats2$^{f/f}$; Yap$^{f/+}$; Taz$^{f/+}$; Shh$^{Cre/+}$; ROSA26$^{mTmG/+}$* (Rescued) mice at 18.5 *days post-coitus* (*dpc*). The EGFP signal from the *ROSA26$^{mTmG}$* allele was induced by *Shh$^{Cre}$*. (d-i) Immunostaining of lung sections from control, mutant, and rescued mice. SCGB1A1 marked club cells, Ac-tubulin (Ac-tub) labeled ciliated cells, SFTPC identified alveolar type 2 (AT2) cells, and T1α characterized alveolar type 1 (AT1) cells. (j, k) Quantification of cell types in the airways and saccules of control, mutant, and rescued mice (n = 3 for each genotype).

All values are mean ± SEM. (***) $p < 0.001$; ns not significant (two-way ANOVA).

Scale bars, 1 mm (a-c), 25 μm (d-f), 100 μm (g-i).

Source data are provided as a Source Data file.

Zhang et al. Supplementary Figure 3

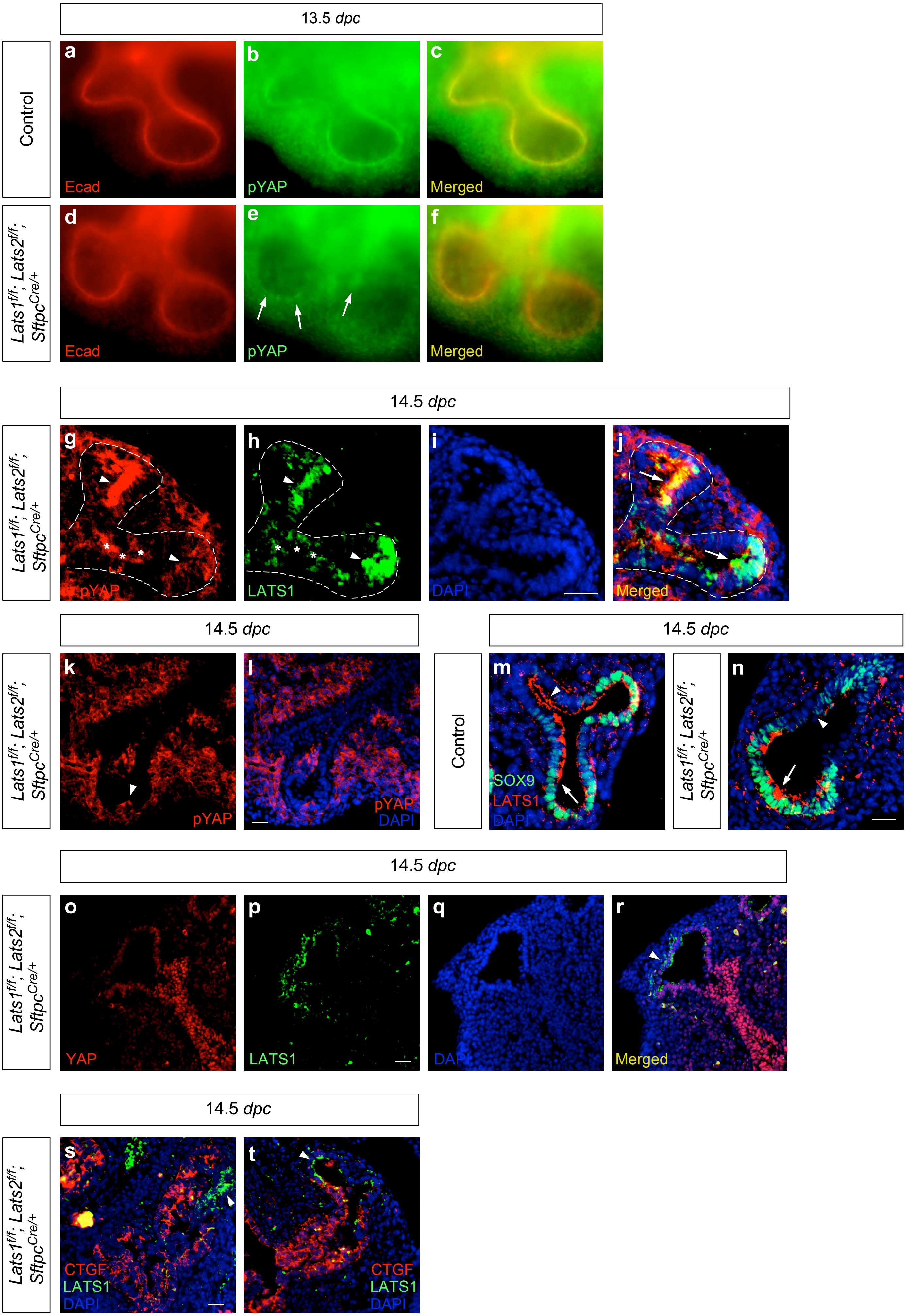

**Supplementary Fig. 3 Regional loss of *Lats1/2* in *Lats1$^{f/f}$; Lats2$^{f/f}$; Sftpc$^{Cre/+}$* lungs is associated with the loss of pYAP and increased YAP**

(a-f) Whole-mount immunostaining of lungs from control and *Lats1$^{f/f}$; Lats2$^{f/f}$; Sftpc$^{Cre/+}$* (*Lats1/2*-mosaic) mice at 13.5 *days post-coitus* (*dpc*). White arrowheads in (e) pointed to residual pYAP signals at the distal tip of *Lats1/2*-mosaic lungs. Ecad (E-cadherin) labeled adherens junctions. (g-l) Immunostaining of lung sections from *Lats1/2*-mosaic mice at 14.5 *dpc*. The white dotted line traced the epithelial layer of the distal lung buds. White arrows pointed to colocalization of residual pYAP and LATS1 signals at the distal tip in (j). The white arrowhead pointed to the residual pYAP (g, k) or LATS1 (h) signal at the distal tip. Signals marked by (*) are from another branch out of the focal place. (m, n) Immunostaining of lung sections from control and *Lats1/2*-mosaic mice at 14.5 *dpc*. White arrows pointed to the SOX9$^{+}$ domain, where the LATS1 signal was also detected. White arrowheads pointed to the SOX2$^{+}$ domain, where the LATS1 signal was present in control lungs but absent in *Lats1/2*-mosaic lungs. (o-r) Immunostaining of lung sections from *Lats1/2*-mosaic mice at 14.5 *dpc*. LATS1 signal (arrowhead) was detected in a subset of the distal SOX9$^{+}$ subdomain. Nuclear YAP signal was significantly reduced in distal epithelial cells carrying the LATS1 signal. By contrast, the nuclear YAP signal was enhanced in proximal epithelial cells lacking the LATS1 signal. (s, t) Immunostaining of lung sections from *Lats1/2*-mosaic mice at 14.5 *dpc*. The CTGF signal was detected in regions where the LATS1 signal was absent. Arrowheads pointed to the residual LATS1 signal at the distal tip.

Scale bars, 25 μm (a-f), 25 μm (g-j), 25 μm (k, l), 25 μm (m, n), 25 μm (o-t).

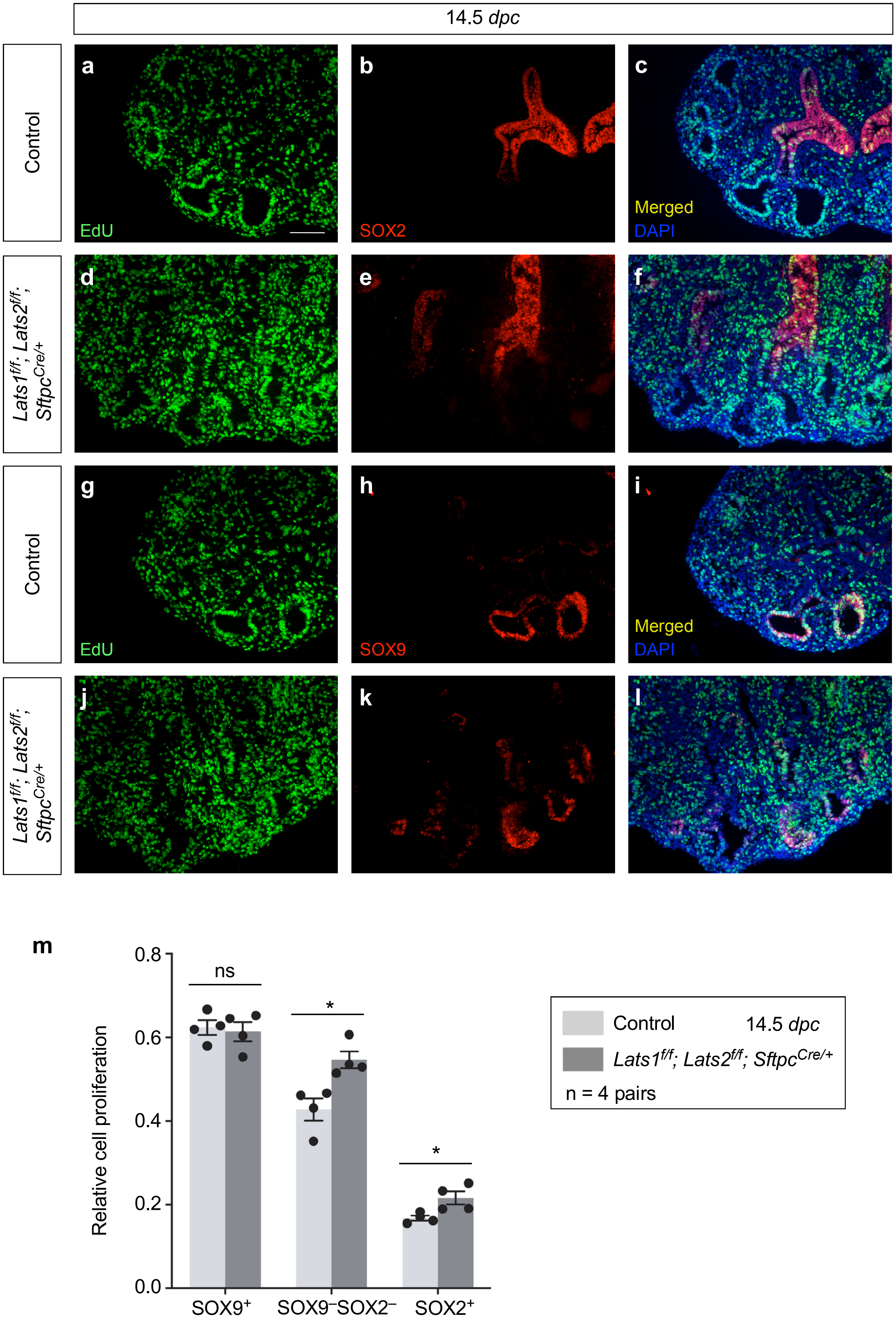
14.5 dpc
Control
Lats1f/f; Lats2f/f; SftpcCre/+
Control
Lats1f/f; Lats2f/f; SftpcCre/+
a
b
c
d
e
f
g
h
i
j
k
l
EdU
SOX2
Merged
DAPI
EdU
SOX9
Merged
DAPI
m
Relative cell proliferation
0.0
0.2
0.4
0.6
0.8
ns
*
*
SOX9+
SOX9−SOX2−
SOX2+
Control
14.5 dpc
Lats1f/f; Lats2f/f; SftpcCre/+
n = 4 pairs

**Supplementary Fig. 4 The proliferation rate is not reduced in the SOX9⁻SOX2⁻ domain of *Lats1$^{f/f}$; Lats2$^{f/f}$; Sftpc$^{Cre/+}$* lungs**

(a-l) Immunostaining of lung sections from control and *Lats1$^{f/f}$; Lats2$^{f/f}$; Sftpc$^{Cre/+}$* (*Lats1/2*-mosaic) mice at 14.5 *days post-coitus* (*dpc*). SOX9 and SOX2 marked the distal and proximal airway epithelium, respectively. EdU labeled proliferating cells. (m) Quantification of the relative cell proliferation in different domains of the lung branches in control and *Lats1/2*-mosaic mice at 14.5 *dpc* (n = 4 pairs), as indicated. The $SOX9^{-}SOX2^{-}$ domain is the intermediate region (transition zone) between the $SOX9^{+}$ and $SOX2^{+}$ domains.

All values are mean ± SEM. (*) $p < 0.05$; ns not significant (two-tailed Student's *t*-test).

Scale bars, 50 μm (a-l).

Source data are provided as a Source Data file.

Zhang et al. Supplementary Figure 5

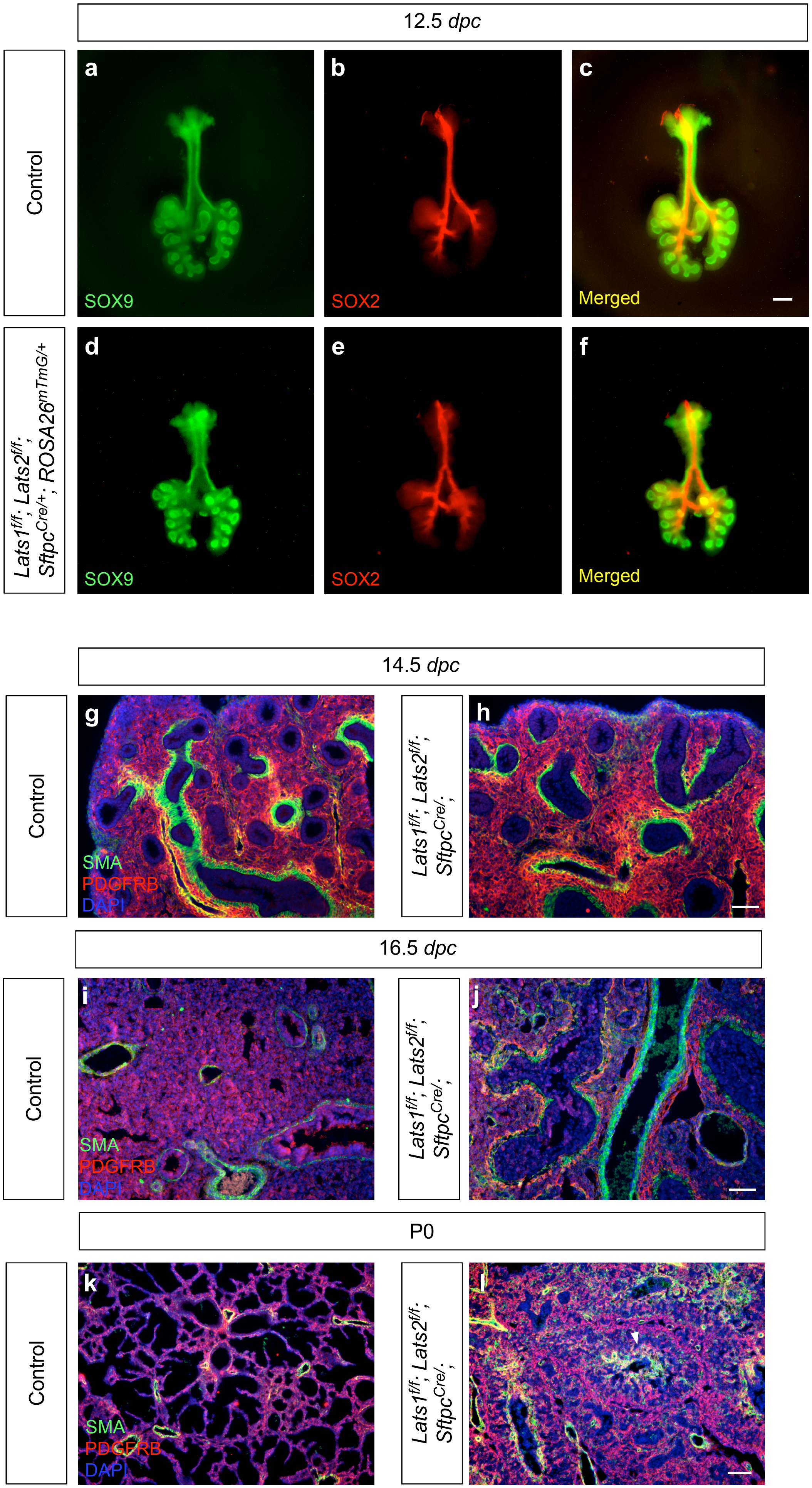

**Supplementary Fig. 5 Early lung branching is not disrupted in *Lats1$^{f/f}$; Lats2$^{f/f}$; Sftpc$^{Cre/+}$* lungs**

(a-f) Whole-mount immunostaining of representative lungs (ventral view) from control and *Lats1$^{f/f}$; Lats2$^{f/f}$; Sftpc$^{Cre/+}$; ROSA26$^{mTmG/+}$* (*Lats1/2*-mosaic) mice at 12.5 *days post-coitus* (*dpc*). SOX9 labeled the distal airway epithelium, while SOX2 labeled the proximal airway epithelium. No apparent difference in the branching pattern was noted between control and *Lats1/2*-mosaic lungs at 12.5 *dpc*. (g-l) Immunostaining of lung sections from control and *Lats1$^{f/f}$; Lats2$^{f/f}$; Sftpc$^{Cre/+}$* (*Lats1/2*-mosaic) mice at 14.5 *dpc*, 16.5 *dpc,* or postnatal (P) day 0. Smooth muscle actin (SMA)-expressing cells include smooth muscle cells and myofibroblasts; PDGFRB-expressing cells include vascular smooth muscle cells and pericytes. The arrowhead in (l) marked a narrow lumen surrounded by thick layers of DAPI-stained epithelial cells. A disorganized epithelium with a similar architecture could be detected in *Lats1$^{f/f}$; Lats2$^{f/f}$; Sftpc$^{Cre/+}$* lungs, shown in (h) of Supplementary Figure 2.

Scale bars, 0.5 mm (a-f), 100 μm (g, h), 100 μm (I, j), 50 μm (k, l).

Zhang et al. Supplementary Figure 6

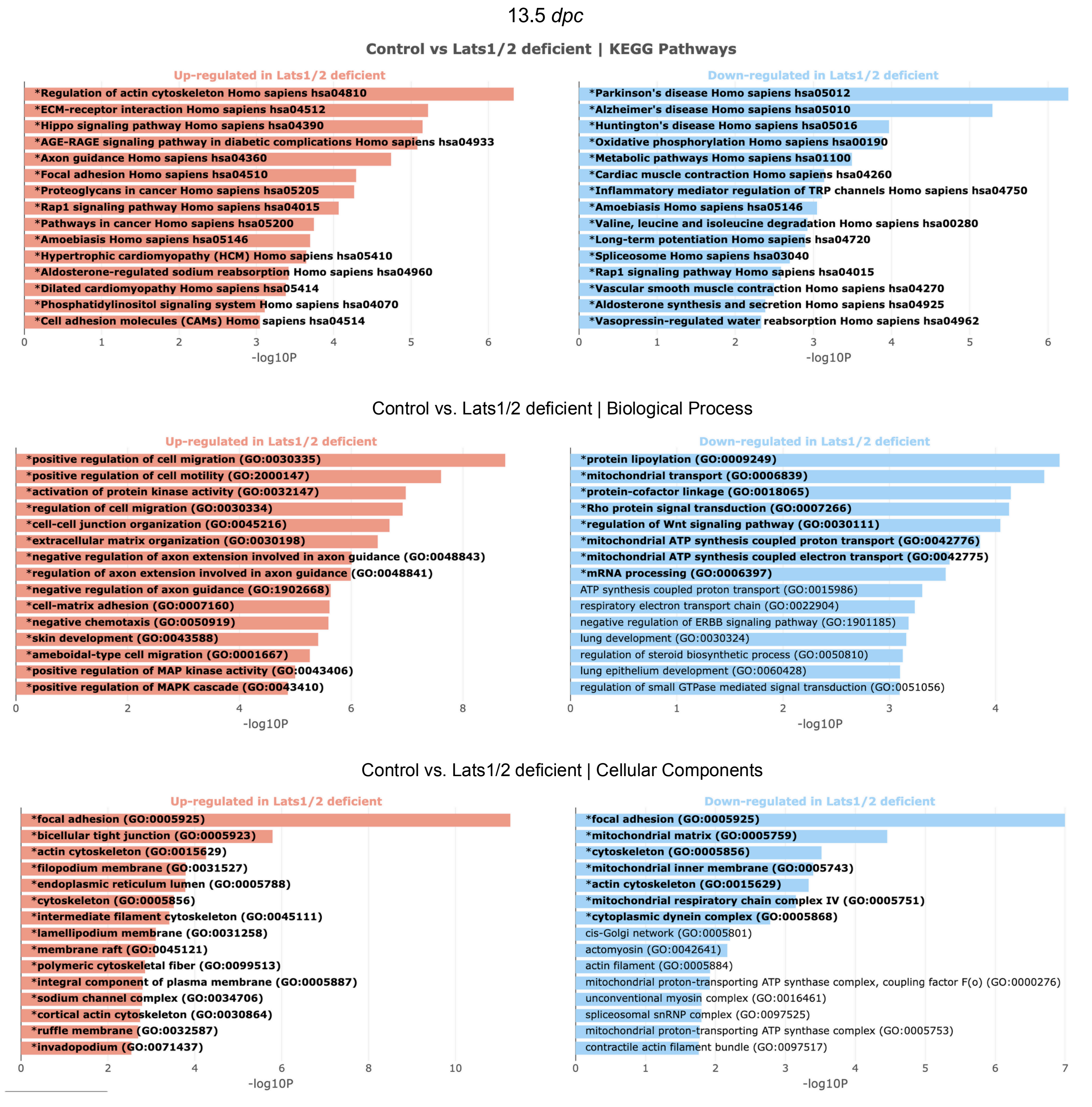

**Supplementary Fig. 6 Pathway analysis of bulk RNA-seq data reveals changes in cellular properties in the absence of *Lats1/2***

Pathway analysis of bulk RNA-seq data from control and $Lats1^{f/f}$; $Lats2^{f/f}$; $Sftpc^{Cre/+}$ mice at 13.5 *days post-coitus* (*dpc*). Pathways regulating focal adhesion, cytoskeleton, cell motility, and cell migration were upregulated in the absence of *Lats1/2*. Each graph shows the top 15 pathways that are enriched. The bold text and asterisk mark pathways with a p-value less than 0.05.

Zhang et al. Supplementary Figure 7

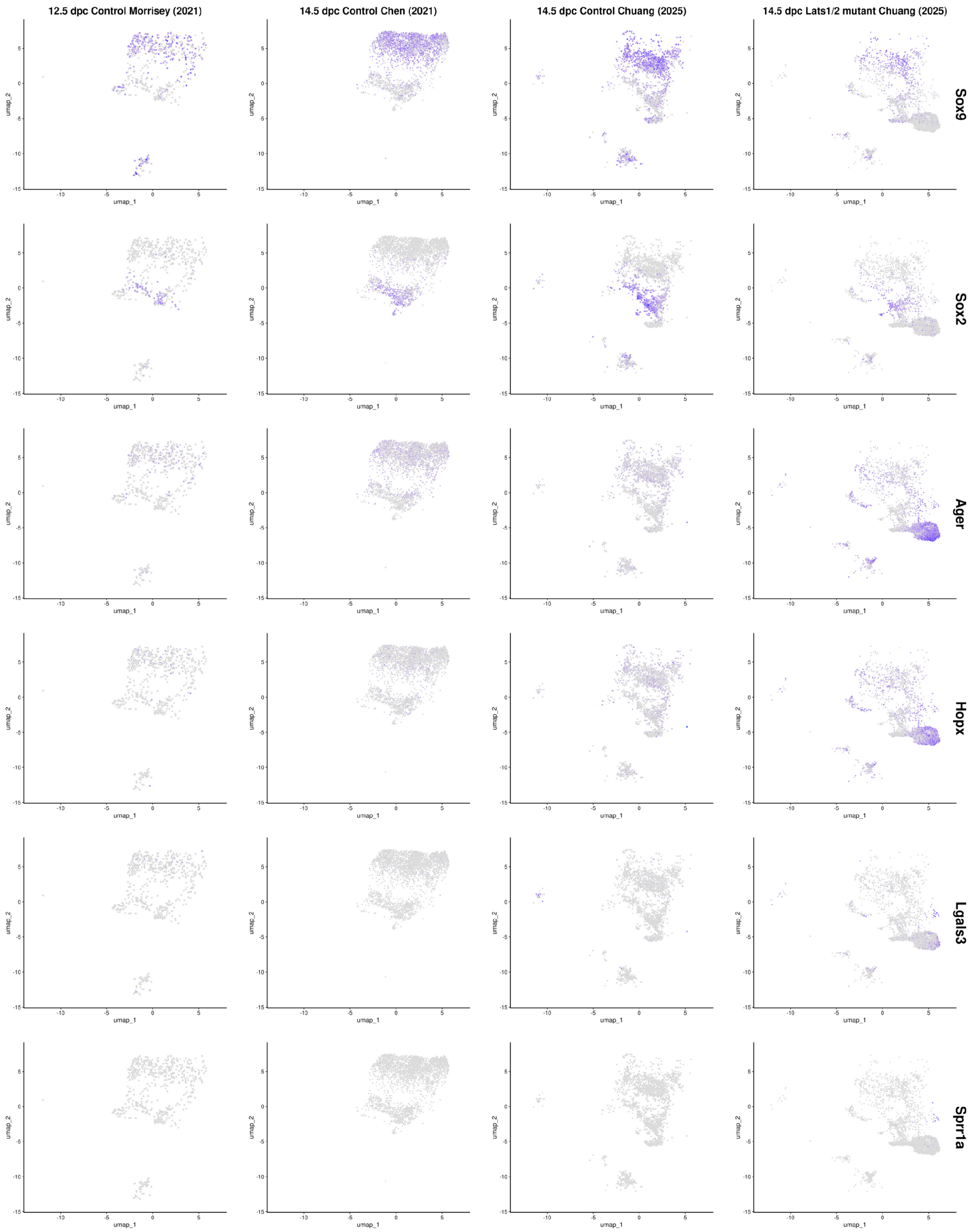

**Supplementary Fig. 7 Re-analysis of published scRNA-seq data at different stages of mouse lung development**

UMAP (uniform manifold approximation and projection) visualization of major cell clusters from scRNA-seq data of control and *Lats1*$^{f/f}$*; Lats2*$^{f/f}$*; Sftpc*$^{Cre/+}$ (*Lats1/2*-mosaic) (Mutant) mouse lungs in this study and those previously published (PMID: 33707239, PMID: 33947861), as specified. Cells that expressed the featured genes were indicated. Expression of *Ager*, *Hopx,* and *Lgals3* was only found in *Lats1/2*-mosaic (Mutant) lungs and not in the other datasets at 14.5 *dpc*.

Zhang et al. Supplementary Figure 8

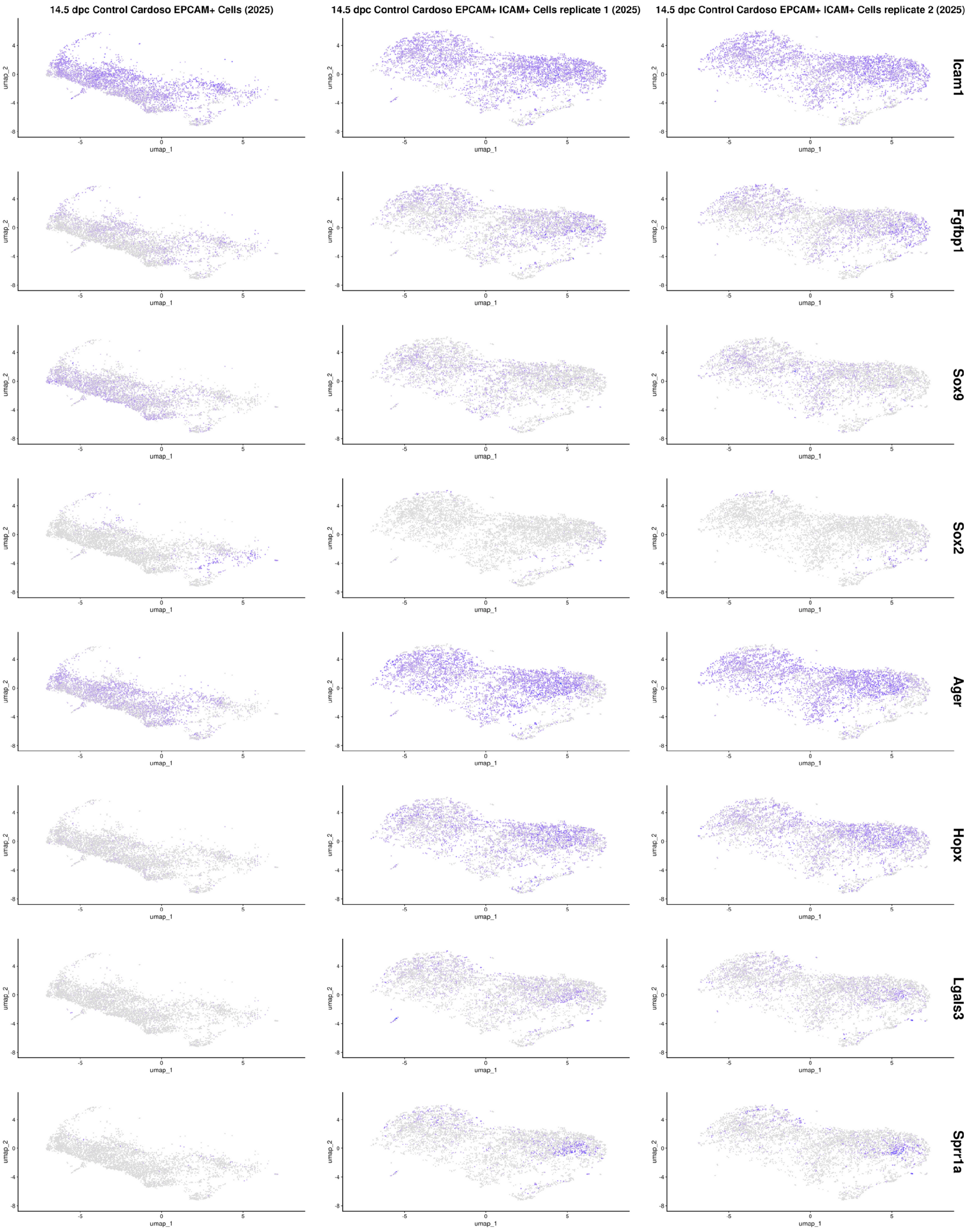

**Supplementary Fig. 8 Re-analysis of published scRNA-seq data at different stages of mouse lung development**

UMAP (uniform manifold approximation and projection) visualization of major cell clusters from scRNA-seq data previously published (PMID: 39667932) as specified. Cells that expressed the featured genes were indicated. Individual samples of published scRNA-seq data were shown. Expression of *Ager*, *Hopx, Lgals3,* and *Sprr1a* was found in two of the three samples at 14.5 *dpc*.

Zhang et al. Supplementary Figure 9

**a**

14.5 *dpc*

Control lungs (Control, CT)

*Lats1$^{f/f}$; Lats2$^{f/f}$; Sftpc$^{Cre/+}$* lungs (*Lats1/2*-mosaic) (Mutant, MT)

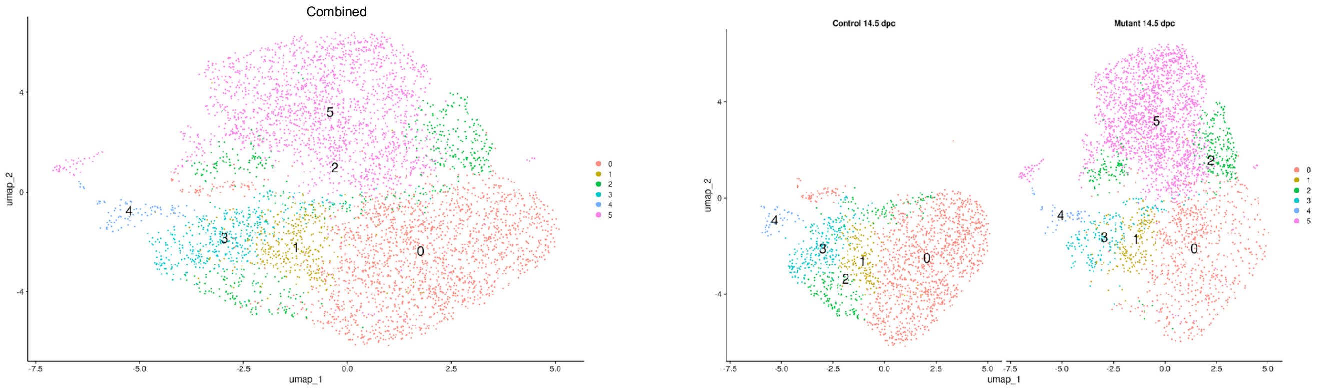


**b**

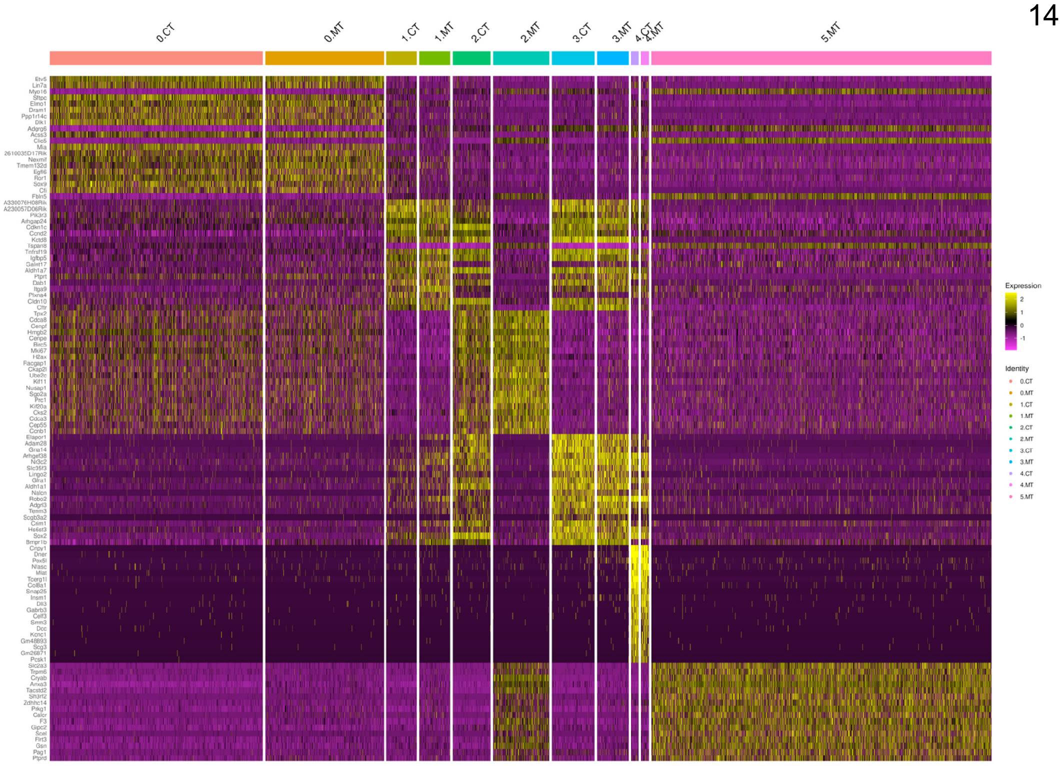


**c**

14.5 *dpc*

| CT | *Sox9*$^{+}$ | *Sox9*$^{+}$ *Sox2*$^{+}$ | *Sox9*$^{–}$ *Sox2*$^{–}$ | *Sox2*$^{+}$ |
|---|---|---|---|---|
| 0 | 74% | 5% | 18% | 3% |
| 1 | 18% | 39% | 12% | 31% |
| 2 | 4% | 41% | 2% | 53% |
| 3 | 2% | 26% | 12% | 60% |
| 4 | 2% | 23% | 5% | 70% |

| MT | *Sox9*$^{+}$ | *Sox9*$^{+}$ *Sox2*$^{+}$ | *Sox9*$^{–}$ *Sox2*$^{–}$ | *Sox2*$^{+}$ |
|---|---|---|---|---|
| 0 | 62% | 6% | 28% | 4% |
| 1 | 9% | 28% | 29% | 34% |
| 2 | 23% | 9% | 45% | 23% |
| 3 | 3% | 18% | 22% | 57% |
| 4 | 7% | 11% | 21% | 61% |
| 5 | 13% | 3% | 67% | 17% |

**Supplementary Fig. 9 Cell clusters in control and *Lats1/2*-mosaic lungs during branching are distinguished by sets of markers**

(a) A combined scRNA-seq analysis of control (CT) and *Lats1*$^{f/f}$; *Lats2*$^{f/f}$; *Sftpc*$^{Cre/+}$ (*Lats1/2*-mosaic) (Mutant, MT) mouse lungs at 14.5 *days post-coitus* (*dpc*). The left panel showed the composite of cell clusters, while the right panel showed the respective cell clusters for control or mutant lungs. (b) Heatmap of cell clusters in control and *Lats1/2*-mosaic lungs. Cluster 2 in control and *Lats1/2*-mosaic lungs shares specific patterns of gene expression (heatmap), but could also be distinguished by other unique sets of gene expression. For instance, clusters 2 and 5 in *Lats1/2*-mosaic lungs share specific patterns of gene expression that are not present in cluster 2 of the control lungs. (c) The % of cells in each cluster denoted by numbers that expressed the indicated marker (or lack thereof) is specified.

Zhang et al. Supplementary Figure 10

14.5 *dpc*

Control lungs (Control, CT)

*Lats1*$^{f/f}$*; Lats2*$^{f/f}$*; Sftpc*$^{Cre/+}$ lungs (*Lats1/2*-mosaic) (Mutant, MT)

**a**

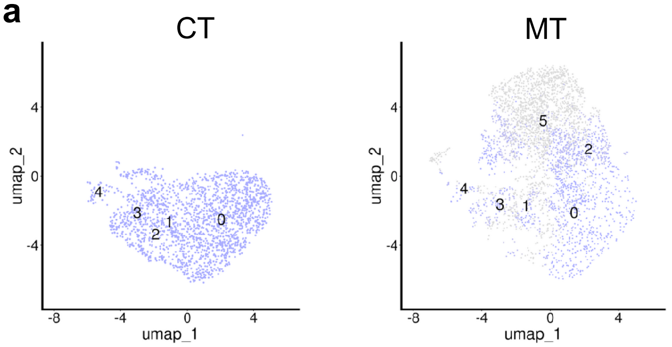


**b**

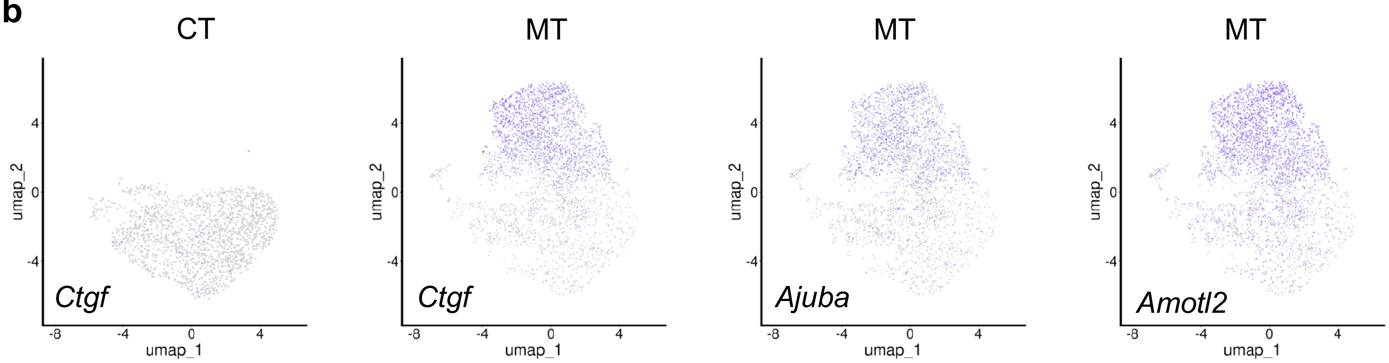


17.5 *dpc*

Control lungs (Control, CT)

*Lats1*$^{f/f}$*; Lats2*$^{f/f}$*; Sftpc*$^{Cre/+}$ lungs (*Lats1/2*-mosaic) (Mutant, MT)

**c**

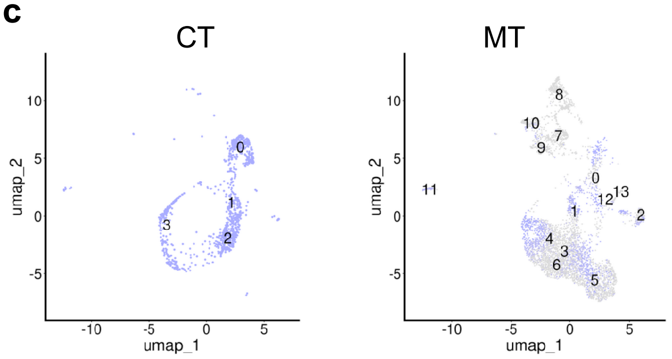


**d**

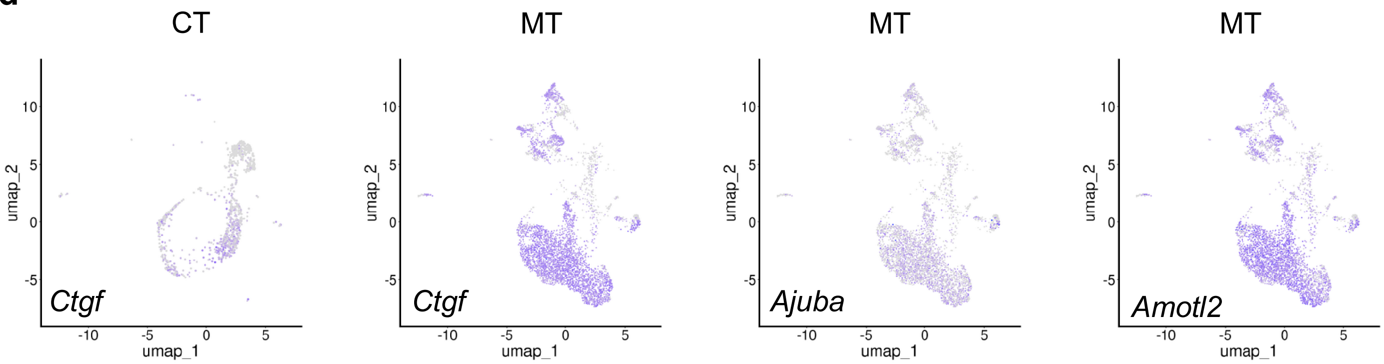

**Supplementary Fig. 10 scRNA-seq analysis of control and *Lats1/2*-mosaic mouse lungs reveals the distribution and trajectory of mutant cells**

(a) UMAP (uniform manifold approximation and projection) visualization of major cell clusters from scRNA-seq data of control (CT) and *Lats1*$^{f/f}$; *Lats2*$^{f/f}$; *Sftpc*$^{Cre/+}$ (*Lats1/2*-mosaic) (Mutant, MT) mouse lungs at 14.5 *days post-coitus* (*dpc*). Cells in the mutant lungs with a transcriptome similar to that in the control lungs were labeled, representing wild-type cells in the mutant lungs at 14.5 *dpc*. Note that the cell clusters and their numbers depicted here correspond to those in Fig. 6. (b) UMAP visualization of representative YAP targets (*e.g.*, *Ctgf*, *Ajuba*, and *Amotl2*) in control and mutant lungs at 14.5 *dpc*. (c) UMAP visualization of the major cell clusters from control and mutant mouse lungs at 17.5 *dpc*. Cells in the mutant lungs with a transcriptome similar to that in the control lungs were labeled, representing wild-type cells in the mutant lungs at 17.5 *dpc*. Note that the cell clusters and their numbers depicted here correspond to those in Fig. 7. (d) UMAP visualization of representative YAP targets (*e.g.*, *Ctgf*, *Ajuba*, and *Amotl2*) in control and mutant mouse lungs at 17.5 *dpc*.

**a**

Control lungs 14.5 *dpc*   Increased in expression ↑   Reduced in expression ↓

| | | |
|---|---|---|
| Cluster 0 — 1 | *Anxa1 Hivep2 Kdm7a Mctp2 Meis2 Mxd4 Mycl Nr3c2 Rarb Rorc Sox2 Tbx1* | ↑ |
| Cluster 0 — 1 | *Brca1 Ccdc25 Celf4 E2f7 E2f8 Etv5 Hirip3 Kif22 Lef1 Nup107 Rbpjl Scx Sp5 Tcf7* | ↓ |
| Cluster 1 — 2 | *Anxa1 Bcl6 Dach1 Hey1 Id4 Kif22 Meis2 Mycl Nr3c2 Ovol1 Six1 Sox2 Sox21 Terf1 Zbtb7c* | ↑ |
| Cluster 1 — 2 | *Creb5 Mycn Zfp521* | ↓ |
| Cluster 2 — 3 | *Npas2 Nr3c2 Trp63* | ↑ |
| Cluster 2 — 3 | *Bcl11b Brca1 Etv5 Foxm1 Hirip3 Kif22* | ↓ |

**b**

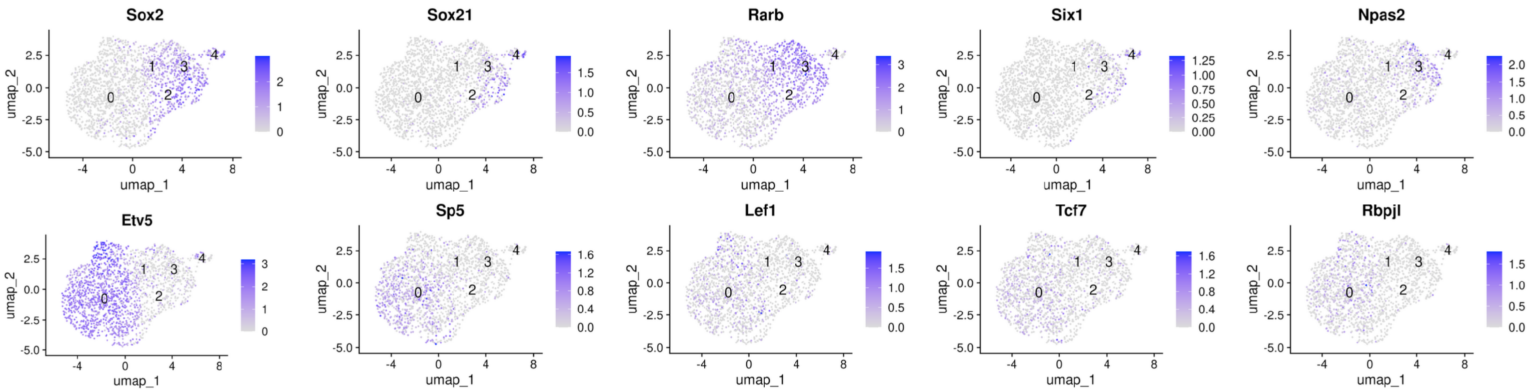


**c**

*Lats1$^{f/f}$; Lats2$^{f/f}$; Sftpc$^{Cre/+}$* lungs 14.5 *dpc*   Increased in expression ↑   Reduced in expression ↓

| | | |
|---|---|---|
| Cluster 0 — 1 | *Ascl1 Bcl6 Egr3 Grhl3 Hivep2 Junb Kdm7a Maf Mctp2 Meis1 Meis2 Mycl Nr3c2 Ovol1 Ppargc1a Prox1 Rarb Sox2 Tbx1* | ↑ |
| Cluster 0 — 1 | *Bcl11b Brca1 Celf4 Dnmt1 Dtl Etv5 Foxm1 Irx1 Lef1 Pold2 Rbpjl Sox9 Sp5 Z2f8 Zfp367* | ↓ |
| Cluster 0 — 2 | *Anxa1 Dmrt1 Elf4 Jdp2 Osr1 Pou6f2 Runx2 Zbtb8b Zbtb32* | ↑ |
| Cluster 0 — 2 | *Etv5 Sox9 Elf5 Lef1 Celf4 Mitf Rbpjl Hivep3 Etv4* | ↓ |
| Cluster 1 — 3 | *Anxa1 Bcl6 Elf3 Fos Id4 Mef2c Meis1 Meis2 Npas2 Ppargc1a Nr3c2 Rorb Sox21* | ↑ |
| Cluster 1 — 3 | *Creb5* | ↓ |
| Cluster 2 — 5 | *Tbx4 Zfp423 Pnox2 Frem1 Bnc2 Meis1 Zeb2 Gli2 Hivep3 Prdm6 Tbx5 Mylk Etv1* | ↑ |
| Cluster 2 — 5 | *Foxm1 Kif22 Mxd3* | ↓ |

**d**

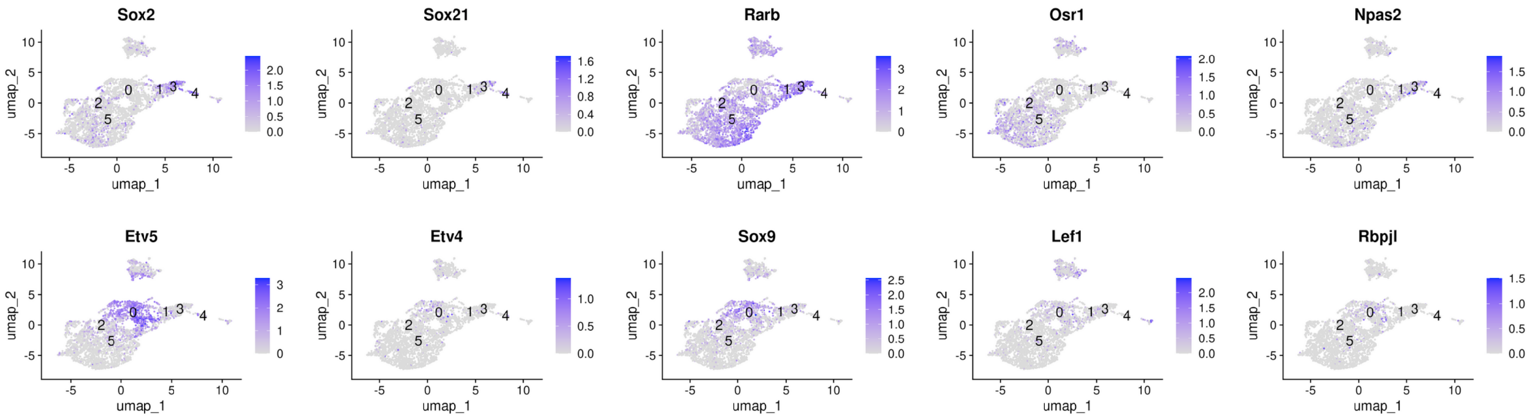

**Supplementary Fig. 11 Transitional cell states from the SOX9$^+$ to SOX2$^+$ state are associated with changes in the expression of transcription factors**

(a) A partial list of transcription factors, the expression of which was either upregulated (blue) or downregulated (red) between two transitional cell states in control mouse lungs at 14.5 *days post-coitus* (*dpc*) as indicated. For instance, genes colored in blue were upregulated while genes colored in red were downregulated when cells transitioned from cluster 0 to 1. (b) UMAP (uniform manifold approximation and projection) visualization of a select group of transcription factors from the partial list for the control mouse lungs. Note that the cell clusters and their numbers depicted here correspond to those in Fig. 6. (c) A partial list of transcription factors, the expression of which was either upregulated (blue) or downregulated (red) between two transitional cell states in *Lats1$^{f/f}$; Lats2$^{f/f}$; Sftpc$^{Cre/+}$* (*Lats1/2*-mosaic) mouse lungs at 14.5 *dpc,* as indicated. (d) UMAP visualization of a select group of transcription factors from the partial list for *Lats1/2*-mosaic mouse lungs. Note that the cell clusters and their numbers depicted here correspond to those in Fig. 6.

Control lungs 14.5 *dpc* Cluster 0 – 1

*Rarb/Meis2/Mxd4/Etv5/E2f7/Sp5/Sox2*

Node type ● DEGs ● TF ● Within top 100 DEGs Correlation direction - Positive - Negative - Undetermined

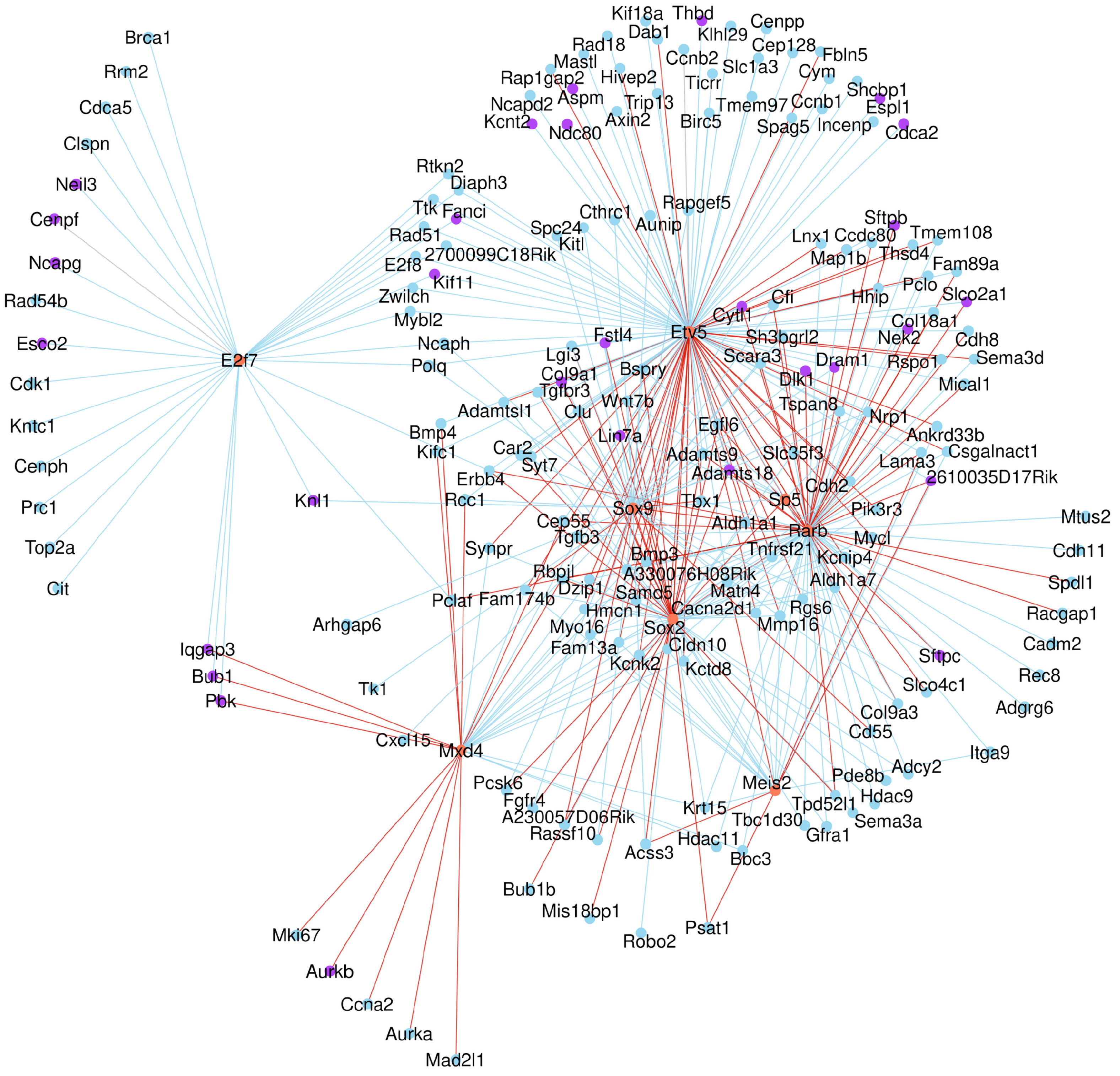

**Supplementary Fig. 12 Transcription factor network for cell state transition during SOX9 to SOX2 transition**

A transcription factor network that centers on transcription factors, *Rarb*, *Meis2*, *Mxd4, Etv5, E2f7, Sp5,* and *Sox2*, which are differentially expressed between cluster 0 and cluster 1 (Fig. 6a, Supplementary Fig. 11a). The potential targets of these transcription factors shown are differentially expressed genes (DEGs) between cluster 0 and cluster 1 (Fig. 6a, Supplementary Fig. 11a). The blue lines indicate positive regulation, while the red lines indicate negative regulation.

Zhang et al. Supplementary Figure 13

**a**

Motifs enriched in $SOX9^{+}$ cells of control lungs at 14.5 *dpc*

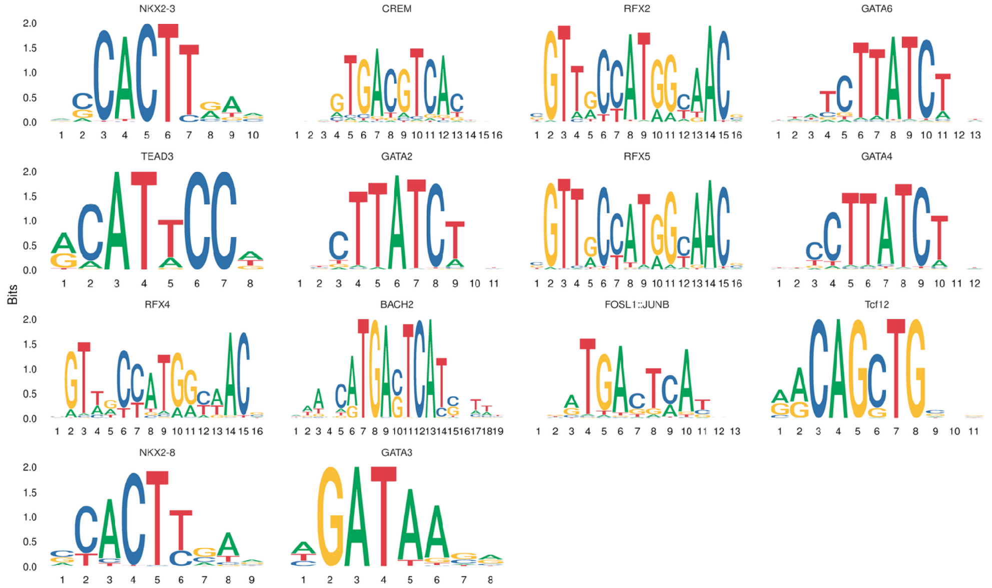


**b**

Motifs enriched in $SOX2^{+}$ cells of control lungs at 14.5 *dpc*

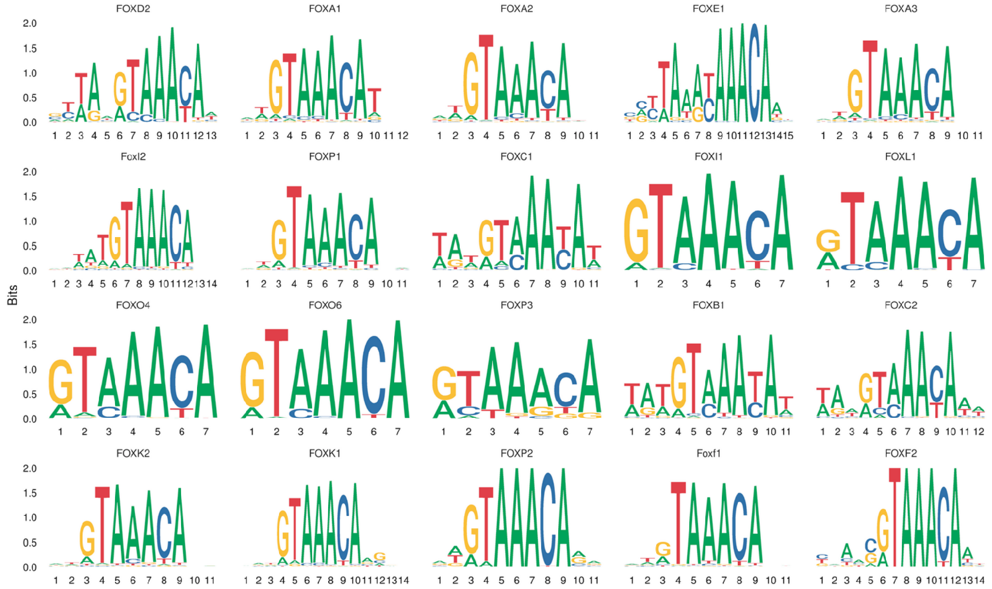


**c**

| DEGs | Motifs | DEGs | Motifs |
|---|---|---|---|
| *Rarb* | TEAD4 | *Etv5* | TEAD4 |
| *Meis2* | TEAD4 | *E2f7* | TEAD4 |
| *Mxd4* | TEAD4 | *Sp5* | TEAD4 |
| *Sox2* | TEAD1 | *Tcf7* | TEAD4 |
| | | *Lef1* | TEAD4 |
| | | *Rbpjl* | TEAD4 |

**Supplementary Fig. 13 Transitional cell states from the SOX9$^+$ to SOX2$^+$ state are associated with changes in open chromatin**

(a) A partial list of motifs identified from multiomics analysis of control mouse lungs at 14.5 *days post-coitus* (*dpc*). These motifs were enriched in SOX9$^+$ cells (relative to SOX2$^+$ cells) within differentially accessible regions (DARs). (b) A partial list of motifs enriched in SOX2$^+$ cells (relative to SOX9$^+$ cells) within DARs. (c) A partial list of differentially expressed genes (DEGs) between the SOX9$^+$ and SOX2$^+$ states that contain the TEAD motif in their regulatory regions.

Zhang et al. Supplementary Figure 14

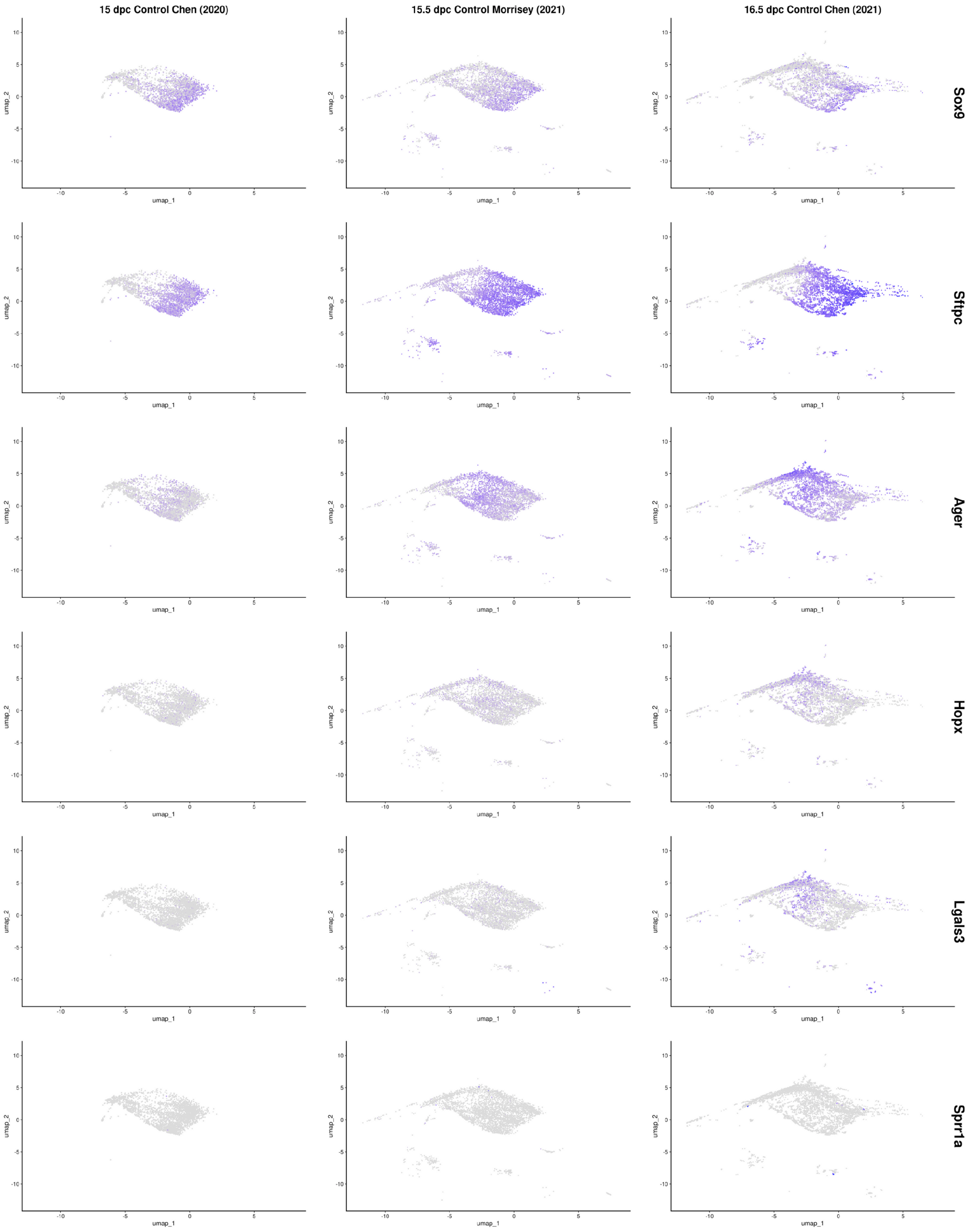

**Supplementary Fig. 14 Re-analysis of published scRNA-seq data at different stages of mouse lung development**

UMAP (uniform manifold approximation and projection) visualization of major cell clusters from scRNA-seq data previously published (PMID: 32414917, PMID: 33707239, PMID: 33947861) as specified. Cells that expressed the featured genes were indicated.

Zhang et al. Supplementary Figure 15

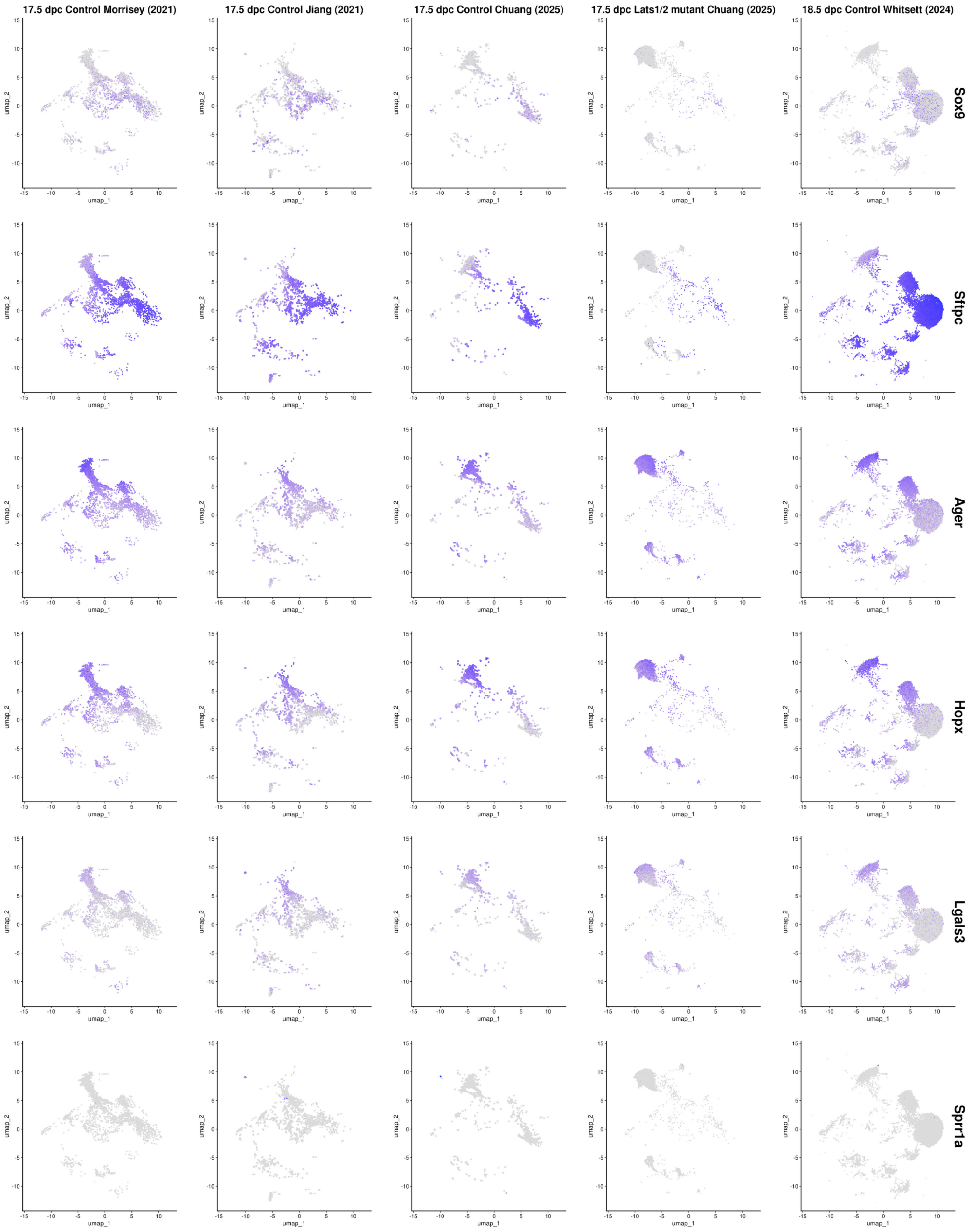

**Supplementary Fig. 15 Re-analysis of published scRNA-seq data at different stages of mouse lung development**

UMAP (uniform manifold approximation and projection) visualization of major cell clusters from scRNA-seq data of control and *Lats1*$^{f/f}$*; Lats2*$^{f/f}$*; Sftpc*$^{Cre/+}$ (*Lats1/2*-mosaic) (Mutant) mouse lungs in this study and those previously published (PMID: 33707239, PMID: 34151224, PMID: 39284798), as specified. Cells that expressed the featured genes were indicated.

Zhang et al. Supplementary Figure 16

**a** 17.5 *dpc*

Control lungs

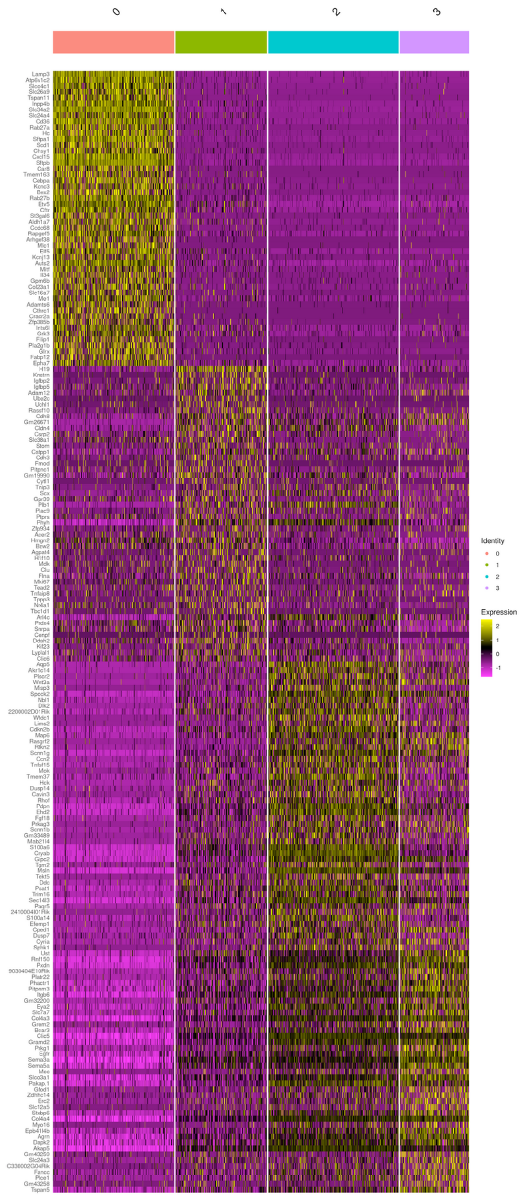


**b** 17.5 *dpc*

*Lats1*$^{f/f}$; *Lats2*$^{f/f}$; *Sftpc*$^{Cre/+}$ lungs

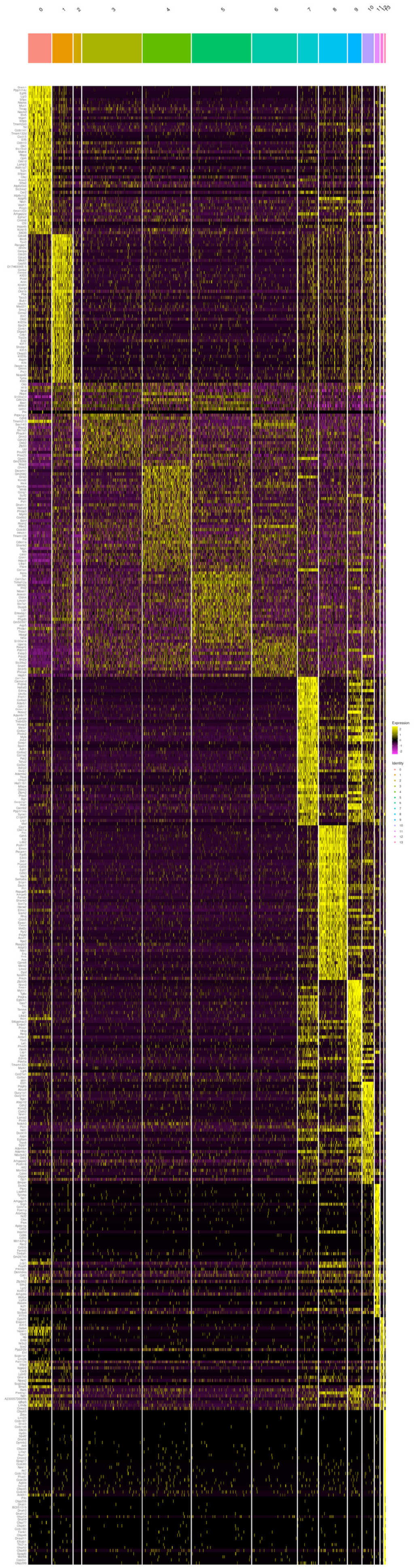

**Supplementary Fig. 16 Cell clusters in control and *Lats1/2*-mosaic mouse lungs during saccMaulation are distinguished by select sets of markers**

(a) Heatmap of cell clusters in control mouse lungs at 17.5 *days post-coitus* (*dpc*). (b) Heatmap of cell clusters in $Lats1^{f/f}$; $Lats2^{f/f}$; $Sftpc^{Cre/+}$ (*Lats1/2*-mosaic) mouse lungs at 17.5 *dpc*. While cell clusters can be identified with specific marker sets, whether they represent functional cell states requires further investigation.

Zhang et al. Supplementary Figure 17

**a**

Control lungs 17.5 *dpc* Increased in expression ↑ Reduced in expression ↓

| | | |
|---|---|---|
| Cluster 0 —— 1 | *Celf5 Cers3 E2f7 Mycn Myrf Pbx4 Rarb Scx Smad9 Sox5 TEAD4 Zbtb8b* | ↑ |
| Cluster 0 —— 1 | *Arg2 Cebpa Creb3l1 Elf5 Etv5 Foxq1 Gpam Mitf Prdm1 Sp5 Tfcp2l1 Zmat4* | ↓ |
| Cluster 1 —— 2 | *Prkaa2 Stat5b TEAD4 Zbtb16* | ↑ |
| Cluster 1 —— 2 | *Dtl Elf5 Etv5 Hmga1 Hmgb2 Magog Nr4a1 Ran Rfc3 Rnaseh2c Sox9 Thap11 Zfp414* | ↓ |
| Cluster 2 —— 3 | *2310011J03Rik Ak6 Borcs8 Csnk2b Id1 Nme1 Prdx5 Rnaseh2c Rpl35 Snapc5 Spag7 Thap11 Zbtb22 Zfp768* | ↑ |

**b**

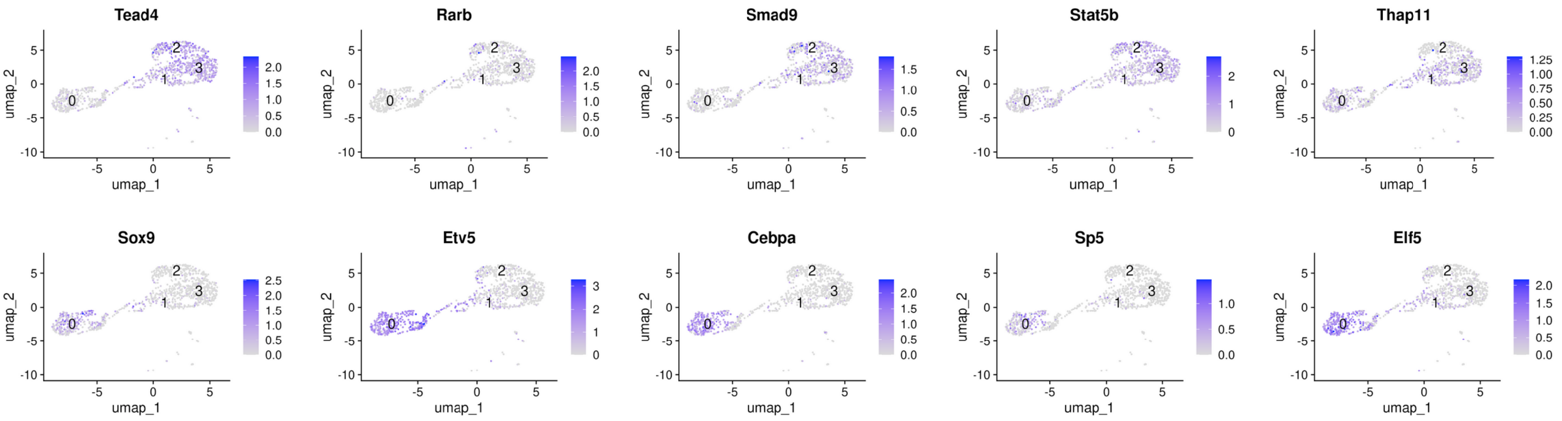


**c**

*Lats1*$^{f/f}$; *Lats2*$^{f/f}$; *Sftpc*$^{Cre/+}$ lungs 17.5 *dpc* Increased in expression ↑ Reduced in expression ↓

| | | |
|---|---|---|
| Cluster 0 —— 1 | *Arid3b Brca1 Dab2 E2f7 E2f8 Elf4 Fhl2 Foxm1 Hirip3 Kif22 Myrf Pbx4 Plagl1 Pou6f2 Runx2 Sox11 Zbtb8b* | ↑ |
| Cluster 0 —— 1 | *Arg2 Ascl2 Bcl6 Cebpa Celf4 Creb3l1 Elf5 Egr2 Epas1 Etv5 Fosb Gpam Hhex Hivep3 Id2 Irf1 Irx2 Irx3 Junb Mitf Nfe212 Nfix Npas2 Nr4a1 Ovol1 Pdrm1 Ppara Ppargc1a Rbpjl Rora Sox9 Sp5 Stat3 Thrb Zfp467 Zmat4* | ↓ |
| Cluster 1 —— 2, 3, 4, 5, 6 | *Hif3a Hivep2 Kdm7a Mxd4 Prkaa2 Tef Zfp949* | ↑ |
| Cluster 1 —— 2, 3, 4, 5, 6 | *Brca1 Dnmt1 Dtl E2f7 E2f8 Elf5 Foxm1 Hirip3 Id1 Kif22 Hmgb2 Mxd3 Mybl2 Timeless* | ↓ |

**d**

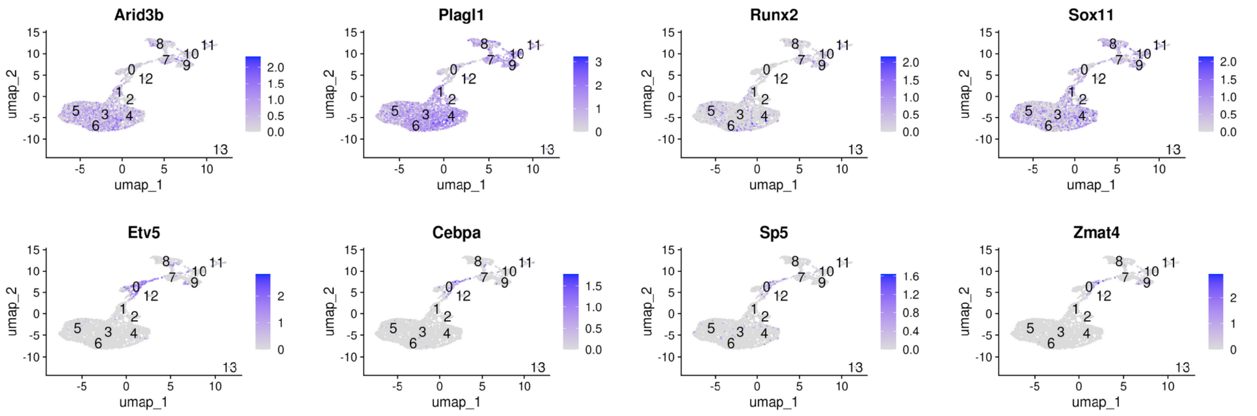

**Supplementary Fig. 17 Transitional cell states from the SOX9$^{+}$ state to the alveolar fate are associated with changes in the expression of transcription factors**

(a) A partial list of transcription factors, the expression of which was either upregulated (blue) or downregulated (red) between two transitional cell states in control mouse lungs at 17.5 *days post-coitus* (*dpc*) as indicated. For instance, genes colored in blue were upregulated while genes colored in red were downregulated when cells transitioned from cluster 0 to 1. (b) UMAP (uniform manifold approximation and projection) visualization of a select group of transcription factors from the partial list for the control mouse lungs. (c) A partial list of transcription factors, the expression of which was either upregulated (blue) or downregulated (red) between two transitional cell states in *Lats1*$^{f/f}$*; Lats2*$^{f/f}$*; Sftpc*$^{Cre/+}$ (*Lats1/2*-mosaic) mouse lungs at 17.5 *dpc,* as indicated. (d) UMAP visualization of a select group of transcription factors from the partial list for *Lats1/2*-mosaic mouse lungs.

Zhang et al. Supplementary Figure 18

Control lungs 17.5 *dpc* Cluster 0 – 1

*Tead4/Rarb/Cebpa/Etv5/Elf5*

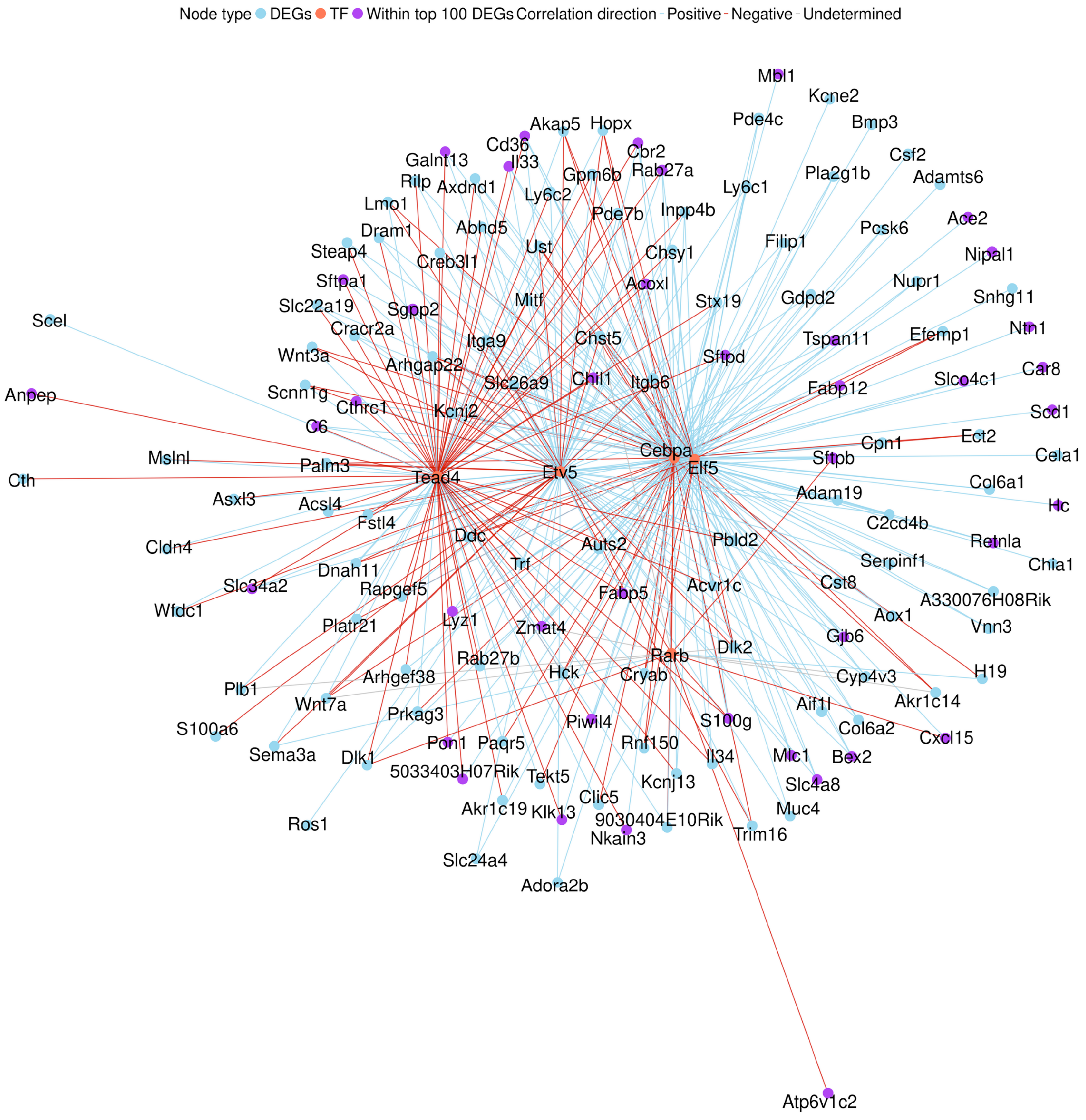

**Supplementary Fig. 18 Transcription factor network for cell state transition during AT2 to AT1 transition**

A transcription factor network that centers on transcription factors, *Tead4*, *Rarb*, *Cebpa, Etv5,* and *Elf5*, which are differentially expressed between cluster 0 and cluster 1 (Fig. 7a, Supplementary Fig. 17a). The potential targets of these transcription factors shown are differentially expressed genes (DEGs) between cluster 0 and cluster 1 (Fig. 7a, Supplementary Fig. 17a). The blue lines indicate positive regulation, while the red lines indicate negative regulation.

**a**

Motifs enriched in AT1 cells of control lungs at 17.5 *dpc*

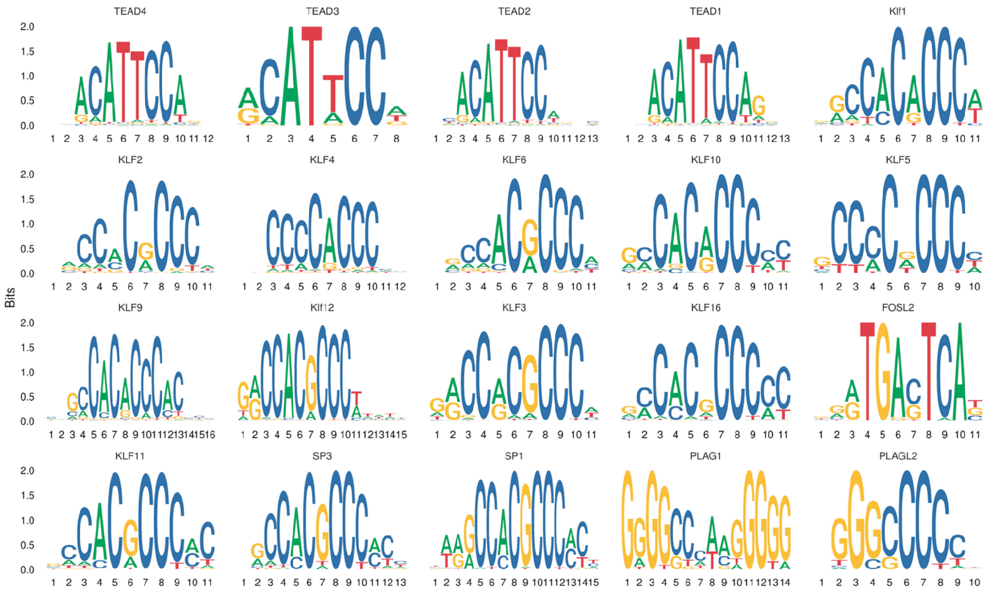


**b**

Motifs enriched in AT2 cells of control lungs at 17.5 *dpc*

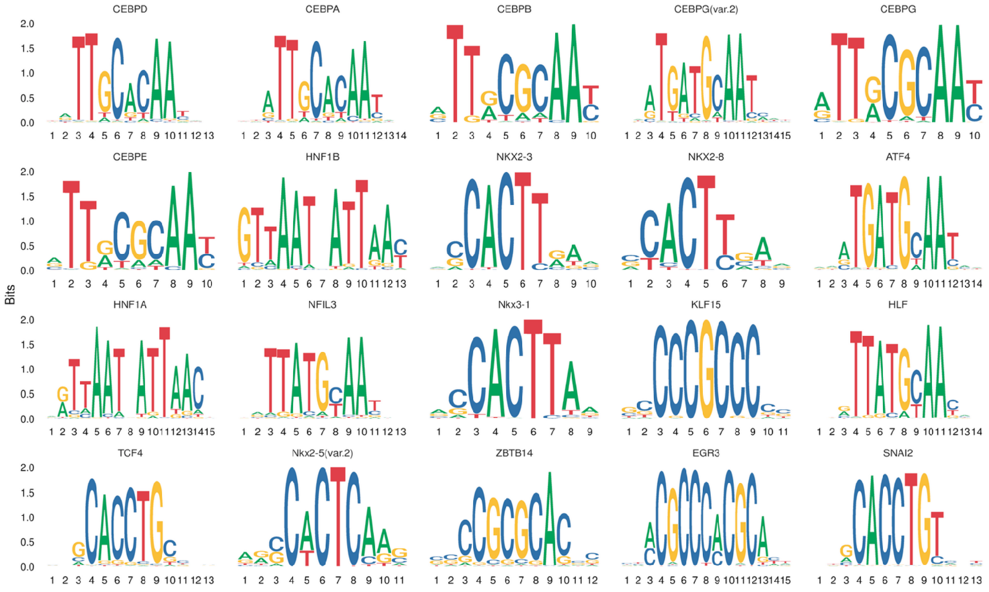


**c**

| DEGs | Motifs |
|---|---|
| *Arid3* | TEAD1 |
| *Klf5* | TEAD4 |
| *Sox11* | TEAD4 |
| *Runx2* | TEAD4 |
| *Plagl1* | TEAD1 |

| DEGs | Motifs |
|---|---|
| *Cebpa* | TEAD4 |
| *Etv5* | TEAD4 |

**Supplementary Fig. 19 Transitional cell states from the SOX9$^{+}$ state to the alveolar fate are associated with changes in open chromatin**

(a) A partial list of motifs identified from multiomics analysis of control mouse lungs at 17.5 *days post-coitus* (*dpc*). These motifs were enriched in AT1 cells (relative to AT2 cells) within differentially accessible regions (DARs). (b) A partial list of motifs enriched in AT2 cells (relative to AT1 cells) within DARs. (c) A partial list of motifs identified in the regulatory regions of differentially expressed genes (DEGs) in AT1 cells (relative to AT2 cells). A partial list of differentially expressed genes (DEGs) between the SOX9$^{+}$ and AT1 states that contain the TEAD motif in their regulatory regions.

Zhang et al. Supplementary Figure 20

**a**

Control mouse lungs 14.5 *dpc*
Cluster 0
Human lungs 5 and 6.86 pcw
Early tip/early stalk

Control mouse lungs 14.5 *dpc*
Cluster 2
Human lungs 5 and 6.86 pcw
Early stalk/early airway progenitor

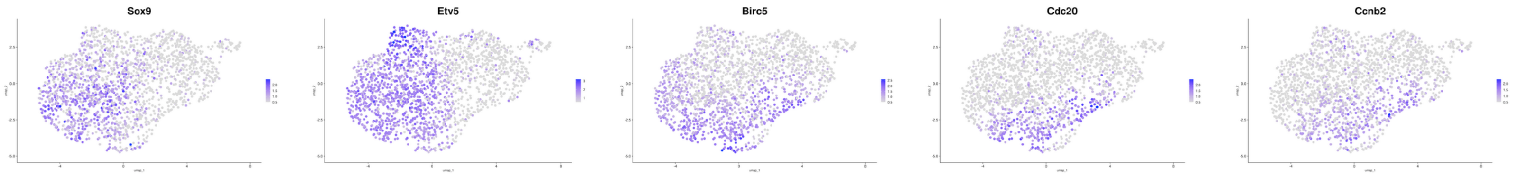


**b**

$Lats1^{f/f}$; $Lats2^{f/f}$; $Sftpc^{Cre/+}$ mouse lungs 14.5 *dpc*
Cluster 0
Human lungs 5 and 6.86 pcw
Early tip/early stalk

$Lats1^{f/f}$; $Lats2^{f/f}$; $Sftpc^{Cre/+}$ mouse lungs 14.5 *dpc*
Cluster 2
Human lungs 5 and 6.86 pcw
Early stalk/early airway progenitor

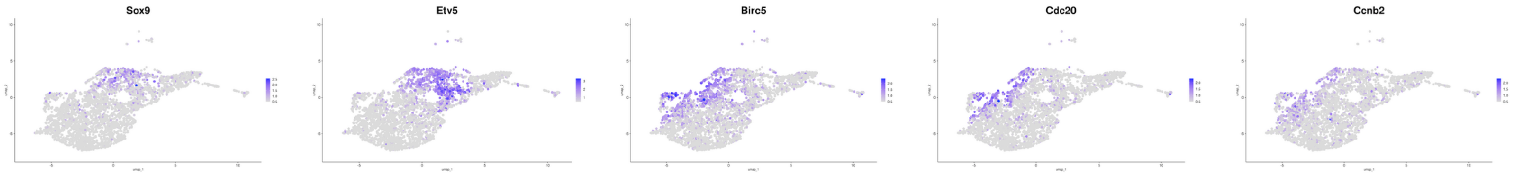


**c**

Control mouse lungs 17.5 *dpc*
Cluster 0
Human lungs 22 pcw
Late tip and others

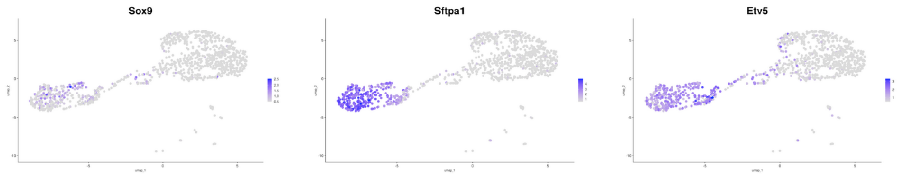


**e**

Control mouse lungs 14.5 *dpc*
Cluster 4
Human lungs neuroendocrine cells

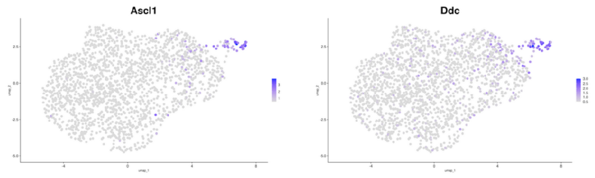


**d**

$Lats1^{f/f}$; $Lats2^{f/f}$; $Sftpc^{Cre/+}$ mouse lungs 17.5 *dpc*
Cluster 0
Human lungs 22 pcw
Late tip and others

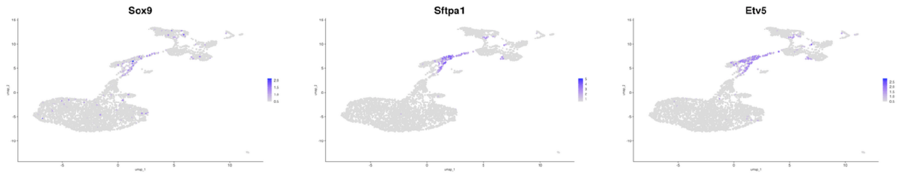


**f**

$Lats1^{f/f}$; $Lats2^{f/f}$; $Sftpc^{Cre/+}$ mouse lungs 14.5 *dpc*
Cluster 4
Human lungs neuroendocrine cells

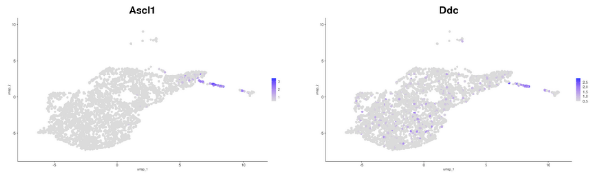

**Supplementary Fig. 20 Cell clusters in the developing mouse lung are connected to distinct regions of the developing human lung**

(a) UMAP (uniform manifold approximation and projection) visualization of representative markers as cells transition from cluster 0 to cluster 2 in control mouse lungs at 14.5 *days post-coitus* (*dpc*) profiled by scRNA-seq. This pattern of gene expression corresponds to the early tip/early stalk and early stalk/early airway progenitor, respectively, in human lungs at 5 and 6.86 post-conception weeks (pcw). (b) UMAPs of representative markers as cells transition from cluster 0 to cluster 2 in *Lats1$^{f/f}$; Lats2$^{f/f}$; Sftpc$^{Cre/+}$* mouse lungs at 14.5 *dpc*. This pattern of gene expression corresponds to the early tip/early stalk and early stalk/early airway progenitor, respectively, in human lungs at 5 and 6.86 pcw. (c) UMAPs of representative markers of cluster 0 in control mouse lungs at 17.5 *dpc*. This pattern of gene expression corresponds to the late tip (and others) of the human lung at 22 pcw. The number of cells in scRNA-seq of human lungs is relatively small, and the conclusion needs to be validated using additional human single-cell data. (d) UMAPs of representative markers of cluster 0 in *Lats1$^{f/f}$; Lats2$^{f/f}$; Sftpc$^{Cre/+}$* mouse lungs at 17.5 *dpc*. This pattern of gene expression corresponds to the late tip of the human lung. (e) UMAPs of representative markers of cluster 4 in control mouse lungs at 14.5 *dpc*. This pattern of gene expression corresponds to neuroendocrine cells in human lungs. (f) UMAPs of representative markers of cluster 4 in *Lats1$^{f/f}$; Lats2$^{f/f}$; Sftpc$^{Cre/+}$* mouse lungs at 14.5 *dpc*. This pattern of gene expression corresponds to neuroendocrine cells in human lungs. In general, conclusions drawn from the correspondence between mouse and human transcriptomes require further analysis using additional markers and cell numbers.

| ECM genes and regulators enriched in $SOX9^+$ cells | ECM genes and regulators enriched in $SOX2^+$ cells |
|---|---|
| **ECM genes** | **ECM genes** |
| *Adamts9* | *Adamtsl1* |
| *Adamts18* | *Dcn* |
| *Ccdc80* | *Fbln5* |
| *Col4a3* | *Lama2* |
| *Col9a1* | *Itgb4* |
| *Col9a3* | *Itih5* |
| *Col15a1* | *Ltbp4* |
| *Col18a1* | *Ptprz1* |
| *Cthrc1* | *Spon1* |
| *Egfl6* | *Thsd4* |
| *Lama3* | |
| *Lamc2* | |
| *Matn4* | |
| *Prelp* | |
| **ECM regulators** | **ECM regulators** |
| *Alpl* | *CCn3* |
| *Bmp1* | *Mmp16* |
| *Fgf9* | *Olfml2a* |
| *Pcsk6* | *Wnt4* |
| *Thbs1* | *Wnt5a* |

**Supplementary Fig. 21 Extracellular matrix (ECM) proteins and their regulators are differentially distributed in SOX9$^+$ and SOX2$^+$ lung cells**

A partial list of ECM genes and their regulators that are enriched in either SOX9$^+$ or SOX2$^+$ mouse lung cells at 14.5 *days post-coitus* (*dpc*). The gene list was derived from differentially expressed genes (DEGs) of scRNA-seq analysis of control mouse lungs at 14.5 *dpc*. Changes in ECM composition could contribute to a difference in the stiffness of the ECM surrounding the SOX9$^+$ (distal) or SOX2$^+$ (proximal) domains in the mouse lung. In this scenario, YAP/TAZ activation is differentially modulated within SOX9$^+$ and SOX2$^+$ domains, or between SOX9$^+$ and SOX2$^+$ domains.

Zhang et al. Supplementary Figure 22

**a**

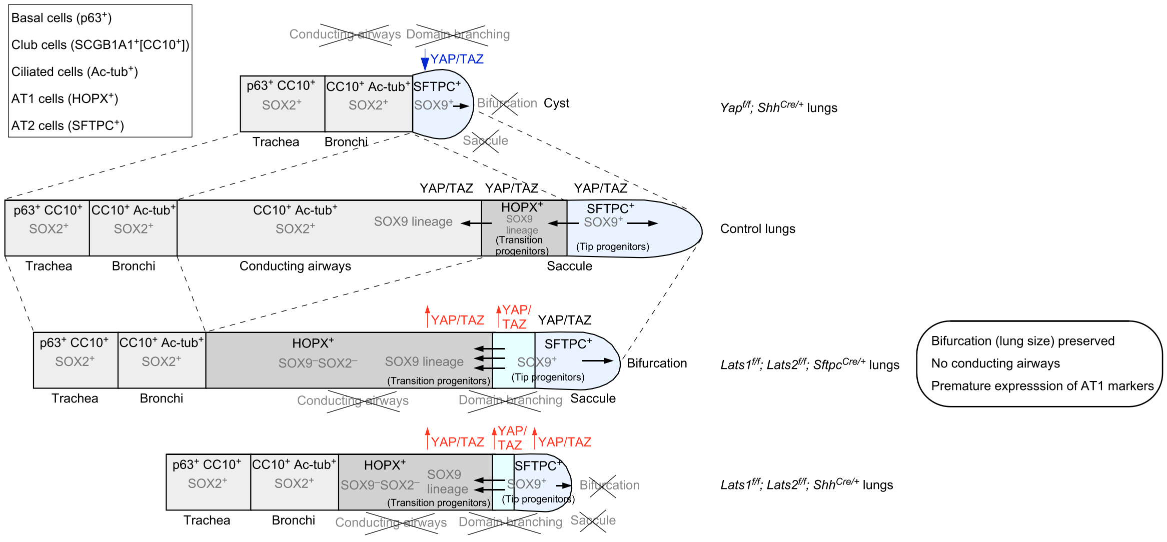


**b**

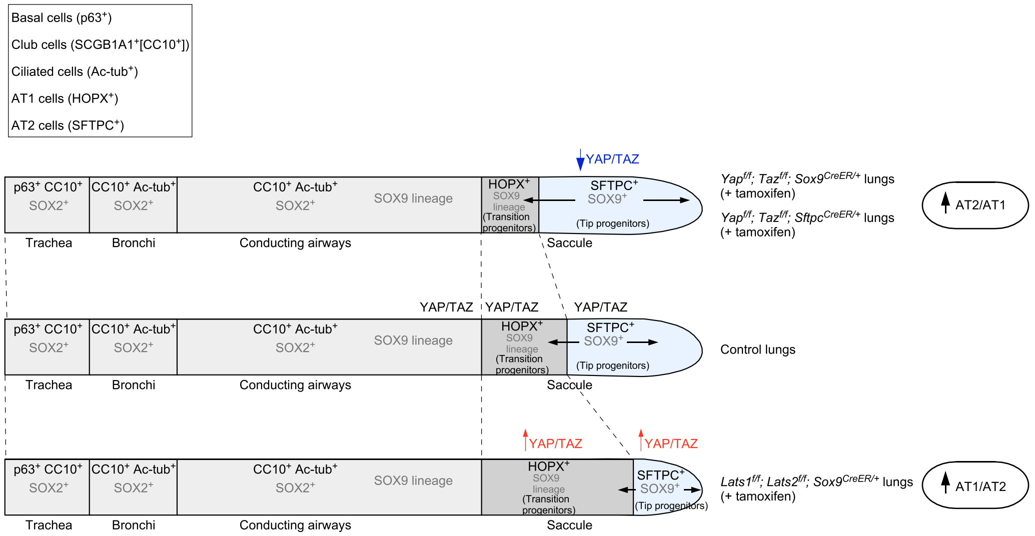

**Supplementary Fig. 22 A model of mouse lung development that is regulated by YAP/TAZ activity**

(a) A model illustrating how YAP/TAZ activity controls lung development, in particular, the SOX9 to SOX2 transition to produce the conducting airways. Lower YAP/TAZ activity (as in $Yap^{f/f}$; $Shh^{Cre/+}$ lungs) fails to maintain $SOX9^{+}$ progenitors and blocks bifurcation and domain branching. Higher YAP/TAZ activity (*e.g.*, loss of *Lats1/2*) promotes premature differentiation of $SOX9^{+}$ progenitors and blocks bifurcation and domain branching. The removal of *Lats1/2* by $Sftpc^{Cre}$ is mosaic (as in $Lats2^{f/f}$; $Sftpc^{Cre/+}$ lungs). Since the distal $SOX9^{+}$ subdomain is spared, bifurcation is unaffected. In contrast, removal of *Lats1/2* by $Shh^{Cre}$ (as in $Lats1^{f/f}$; $Lats2^{f/f}$; $Shh^{Cre/+}$ lungs) affects both domain branching and bifurcation. As a result, YAP/TAZ is depicted to affect both domain branching and bifurcation. (b) A model of how YAP/TAZ activity controls lung development, in particular, the production of AT1 and AT2 cells to produce the alveoli. Lower YAP/TAZ activity (*e.g.*, loss of *Yap/Taz* induced by tamoxifen as in $Yap^{f/f}$; $Taz^{f/f}$; $Sox9^{CreER/+}$ and $Yap^{f/f}$; $Taz^{f/f}$; $Sftpc^{CreER/+}$ lungs) increases the AT2 to AT1 ratio. Higher YAP/TAZ activity (*e.g.*, loss of *Lats1/2* induced by tamoxifen as in $Lats1^{f/f}$; $Lats2^{f/f}$; $Sox9^{CreER/+}$ lungs) promotes the differentiation of AT1 cells and increases the AT1 to AT2 ratio.
